# Genomic patterns of malignant peripheral nerve sheath tumour (MPNST) evolution correlate with clinical outcome and are detectable in cell-free DNA

**DOI:** 10.1101/2022.05.03.490481

**Authors:** I Cortes-Ciriano, CD Steele, K Piculell, A Al-Ibraheemi, V Eulo, MM Bui, A Chatzipli, BC Dickson, DC Borcherding, A Feber, A Galor, J Hart, KB Jones, JT Jordan, RH Kim, D Lindsay, C Miller, Y Nishida, P Proszek, J Serrano, RT Sundby, JJ Szymanski, NJ Ullrich, D Viskochil, X Wang, Genomics of MPNST (GeM) Consortium, M Snuderl, PJ Park, AM Flanagan, AC Hirbe, N Pillay, DT Miller

**Author notes:** Correspondence and requests for materials should be addressed to David T. Miller (MD, PhD). Contributed equally to leading the consortium (I Cortes-Ciriano, AM Flanagan, AC Hirbe, N Pillay, DT Miller).

## Abstract

Malignant peripheral nerve sheath tumour (MPNST) is an aggressive soft-tissue sarcoma that arises in peripheral nerves. MPNST occurs either sporadically or in people with neurofibromatosis type 1 (NF1), a common cancer predisposition syndrome caused by germline pathogenic variants in *NF1*. Although MPNST is the most common cause of death and morbidity for individuals with NF1, the molecular underpinnings of MPNST pathogenesis remain unclear. Here, we report the analysis of whole-genome sequencing, multi-regional exome sequencing, transcriptomic and methylation profiling data for 95 MPNSTs and precursor lesions (64 NF1-related; 31 sporadic) from 77 individuals. Early events in tumour evolution include biallelic inactivation of *NF1* followed by inactivation of *CDKN2A* and in some cases also *TP53* and polycomb repressive complex 2 (PRC2) genes. Subsequently, both sporadic and NF1-related MPNSTs acquire a high burden of somatic copy number alterations (SCNAs). Our analysis revealed distinct pathways of tumour evolution and immune infiltration associated with inactivation of PRC2 genes and H3K27 trimethylation (H3K27me3) status. Tumours with loss of H3K27me3 evolve through extensive chromosomal losses with retention of chromosome 8 heterozygosity followed by whole genome doubling and chromosome 8 amplification. These tumours show lower levels of immune cell infiltration with low cytotoxic activity and low expression of immune checkpoints. In contrast, tumours with retention of H3K27me3 evolve through extensive genomic instability in the absence of recurrent alterations and exhibit an immune cell-rich phenotype. Specific SCNAs detected in both tumour samples and cell-free DNA (cfDNA) act as a surrogate for loss of H3K27me3 and immune infiltration, and predict prognosis. Our results suggest that SCNA profiling of tumour or cfDNA could serve as a biomarker for early diagnosis and to stratify patients into prognostic and treatment-related subgroups.

## Main

Malignant peripheral nerve sheath tumour (MPNST) is an aggressive soft-tissue sarcoma of peripheral nerves that arises both sporadically and in 8-13% of individuals with neurofibromatosis type 1 (NF1)^1,2^. NF1 is a common cancer predisposition disorder affecting 1:2500 individuals worldwide caused by germline heterozygous loss-of-function pathogenic variants in the *NF1* gene^3,4^, which encodes for neurofibromin, a Ras GTPase-activating protein^5,6^. Activation of the Ras pathway is a driver event in many sporadic cancers in the general population, including up to 30% of melanomas and breast cancers, and nearly 25% of acute myeloid leukemias and glioblastomas, among others^7^. Therefore, understanding the development of cancer as a result of *NF1* loss is not only essential for improving prognostication and therapies for MPNST, but can also help to gain a deeper understanding of *NF1*-related cancer biology.

In individuals with NF1, germline inactivation of *NF1* facilitates the development of multiple tumour types, including histologically benign plexiform neurofibromas (PNs). PNs arise through loss of heterozygosity (LOH) for *NF1* in the Schwann cell lineage^8^, and may develop anywhere in the peripheral nervous system. A subset of tumours subsequently undergoes genomic transformation with progression to MPNST, which is the major cause of morbidity and mortality for individuals with NF1^9^. Malignant transformation has been attributed to genetic alterations, including *NF1* inactivation followed by early loss of *CDKN2A*^10,11^ and, in some cases, PRC2 inactivation as a result of somatic mutations in *SUZ12*, *EED* or *EZH2*^10,12,13^. MPNSTs with PRC2 inactivation undergo loss of trimethylation at lysine 27 of histone H3 (H3K27me3)^14^, which has been associated with a worse prognosis^15,16^. However, these events are not found across all MPNSTs, and variability in histological features adds to the challenge of accurately classifying these tumours, thereby limiting clinical decision-making and therapeutic development.

Previous genomic analyses of MPNSTs were based on small sample size studies and limited pathology characterisation^12,17–20^. As a result, the timing and clinical relevance of the genomic alterations underpinning MPNST development remain poorly understood. To overcome these limitations, we established the international Genomics of MPNST (GeM) consortium with the goal of analysing a large collection of MPNSTs with detailed clinical and pathological characterisation at a multi-omic level^21^. Since H3K27me3 has been shown to correlate with clinical outcome, we sought to better understand the molecular events and downstream functional consequences of PRC2 inactivation in MPNST development. In doing so, we anticipated several opportunities: improving diagnostic accuracy, refining clinical prognosis, identifying potentially effective therapeutic intervention, and adding to the understanding of tumorigenesis for cancers related to *NF1* loss. Here, we describe the genomic landscape and key evolutionary events in MPNST development and progression, and provide evidence that this genomic landscape can be used clinically to identify subgroups of patients with varying prognoses.

## Results

### Tumour sample collection, characterisation and stratification

Through international multi-institutional collaboration, we collected and analysed 95 fresh-frozen tumours (64 NF1-related; 31 sporadic) and matched blood samples from 77 individuals (50 with an NF1 clinical phenotype and 27 sporadic) using whole-genome sequencing (WGS), RNA-sequencing and whole-genome DNA methylation arrays (Fig. 1a and Supplementary Table 1). This collection, which is approximately 10 times larger than any prior WGS or multi-omic studies of MPNSTs^12,17^, includes 85 high-grade MPNSTs, 5 low-grade MPNSTs, 3 atypical neurofibromatous neoplasms of uncertain biologic potential (ANNUBPs) and 2 neurofibromas. Cases were selected based on a pathological diagnosis of MPNST or precursor lesion^9^, availability of clinical information, H3K27me3 immunoreactivity data, and availability of material for multi-omic analysis (Methods). Using the WGS data, we identified an *NF1* germline pathogenic variant in all 50 individuals with an NF1 clinical phenotype (8 with pathogenic microdeletions and complex SVs undetectable by exome sequencing and 2 with deep intronic mutations that generated cryptic exons; Supplementary Table 1, Supplementary Figs. 1-3, and Methods). Next, we confirmed biallelic inactivation of *NF1* in 56/64 (88%) tumour samples arising within the tumours of individuals with NF1. *NF1* inactivation in 49 of the 56 (87.5%) tumours occurred through the loss of the wild-type allele, and biallelic inactivation was identifiable more often in tumours from people with NF1 as compared to sporadic tumours, 10/31 (32%, *P*<0.01, Fisher’s exact test; Supplementary Table 1). Four pathologists performed a consensus classification by assigning the tumours to 4 categories based on the presence or absence of germline *NF1* pathogenic variants (germline vs sporadic) and conventional and non-conventional histopathological features of MPNST (Supplementary Fig. 4). In addition, tumours were classified according to the retention or loss of H3K27me3 using immunohistochemistry^9^ (H3K27me3 retained vs H3K27me3 loss; Methods). Diagnosis using previously published machine learning classifiers of sarcomas based on genome-wide DNA methylation patterns^22^ was only informative for 48/95 tumours (51%), highlighting the need for combining pathology and genomics data for accurate diagnosis (Supplementary Fig. 5).

**Figure 1.**
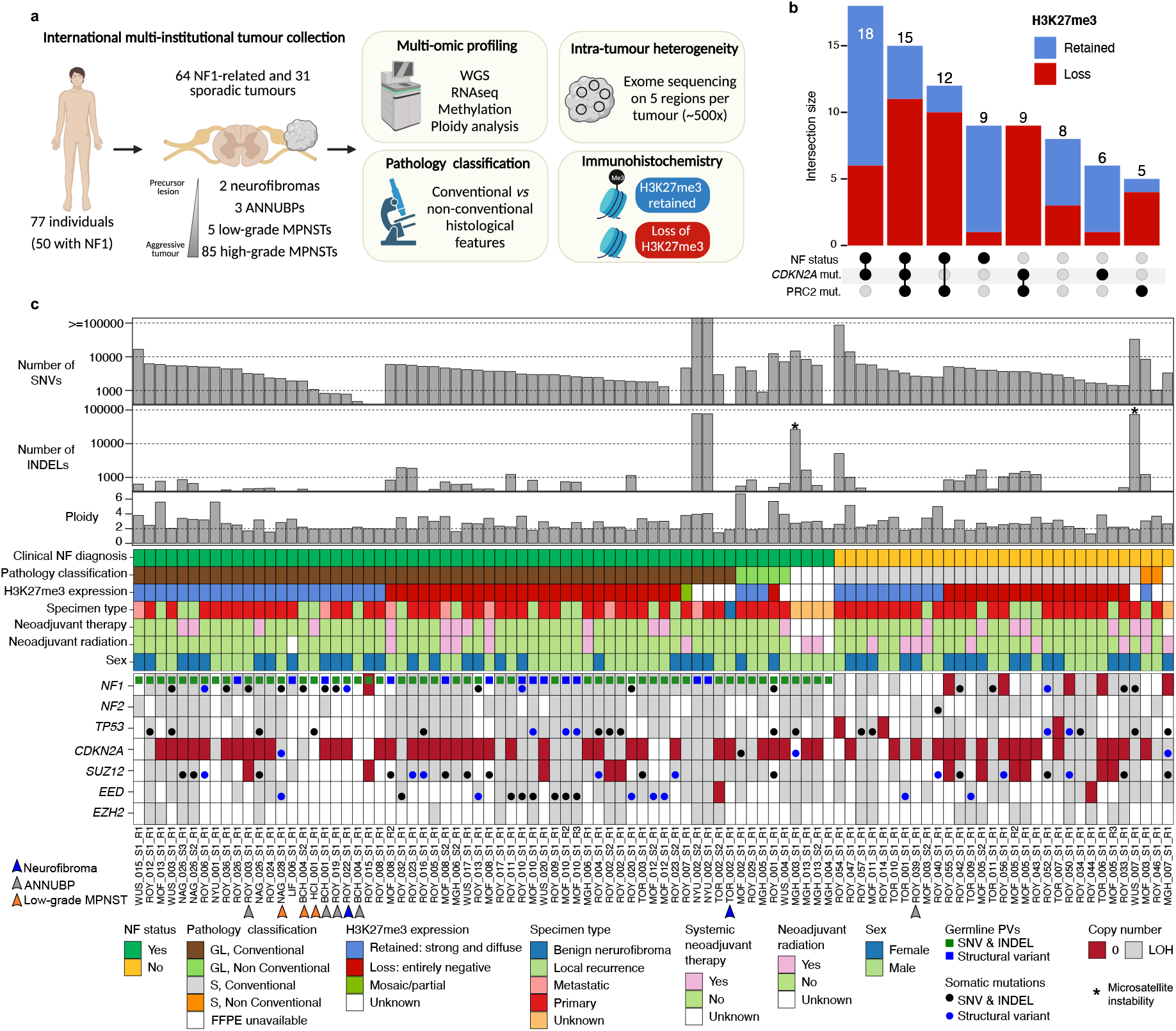
Genomic and pathology classification of MPNSTs. **a,** Overview of the experimental design and technologies utilized in this study. **b**, Relationship between NF status, biallelic inactivation of *CDKN2A* and the PRC2 complex. The numbers on top of the bars indicate the total number of tumours in each group. **c**, Clinical information, histopathological features and mutations in driver genes in the GeM cohort. Precursor lesions are marked by arrowheads. Sample labels not marked by an arrowhead correspond to high-grade MPNSTs. ANNUBP, atypical neurofibromatous neoplasms of uncertain biologic potential; CN, copy number; GL, germline; S, sporadic.

### Somatic genomic landscape of MPNSTs

To identify drivers of MPNST development, we searched for genes harbouring recurrent loss or gain-of-function mutations (Methods). We detected somatic biallelic inactivation of *CDKN2A* in 40/64 (63%) NF1-related MPNSTs and in 17/31 (55%) sporadic cases (Fig. 1b-c), primarily caused by non-recurrent SVs and complex rearrangements of the *CDKN2A* locus, which in some cases led to the concomitant inactivation of *NF1* and the loss of nearby genes (e.g., *MTAP* in 25% of tumours; Supplementary Fig. 6). We identified loss of H3K27me3 in 45/82 (55%) of MPNSTs with immunohistochemistry data available (Fig. 1b-c). *SUZ12* was the gene from the PRC2 complex with the highest frequency of biallelic inactivation (27/95 tumours with WGS data, 28%), followed by *EED* (16/95, 17%). Biallelic inactivation of *SUZ12* and *EED* were mutually exclusive events (Fig. 1c). Furthermore, the loss of H3K27me3 was not associated with hypermethylation of these genes (Supplementary Fig. 7). Double-hit mutations were found in *TP53* in 20/95 (21%) tumours, and replicative immortality was primarily driven by *TERT* overexpression (Supplementary Fig. 8). We did not detect recurrent gene fusions or genes recurrently mutated besides *NF1*, *CDKN2A*, PRC2 complex genes and *TP53* (Fig. 1c and Methods), suggesting that these are the main drivers of MPNST tumorigenesis.

We performed *de novo* mutational signature analysis of single-base substitutions (SBS) and small indels. Similar mutational processes were identified in MPNSTs irrespective of H3K27me3 status (Extended Data Fig. 1a). Mutational signature SBS5, a clock-like signature found in human cancers^23^ and histologically normal tissues^24^, was the predominant signature in both groups, contributing a median of 58% of SNVs. The burden of mutations and SVs was comparable between both groups, with the exception of four hypermutated cases, two of which showed microsatellite instability (Fig. 1c) (Extended Data Fig. 1b-e). SV signature analysis identified five distinct signatures previously detected in undifferentiated soft tissue sarcomas and breast carcinomas^25,26^ (Extended Data Fig. 1f-g), which had comparable activity in both groups (Extended Data Fig. 1a). The frequency of chromothripsis, which was detected in 47/95 (49%) of tumours, was also comparable between both groups (19/37 and 22/45 of tumours with retention and loss of H3K27me3, respectively; *P* > 0.05, Fisher’s exact test). Whole-genome doubling (WGD) events, which were validated using image cytometry analysis (Methods), were detected in 22/37 (59%) of tumours with H3K27me3 retained and in 33/45 (73%) of tumours with H3K27me3 loss (*P* > 0.05, Fisher’s exact test). Together, these results indicate that MPNSTs are characterised by a high level of genomic instability irrespective of H3K27me3 status.

### Molecular subgroups correlate with clinical outcome and immune infiltration

Next, we investigated transcriptomic differences between MPNSTs with different H3K27me3 status. Unbiased clustering based on transcriptomic and genome-wide methylation data confirmed that tumours stratified into two distinct groups, largely corresponding to the retention and loss of H3K27me3 (Methods and Extended Data Fig. 2a). Cluster assignments were consistent across data types (Extended Data Fig. 2a). Classification by H3K27me3 immunoreactivity was associated with overall survival in NF1 individuals, but not in sporadic MPNSTs (Extended Data Fig. 2b-c). No differences in overall survival were observed when stratifying tumours on the basis of conventional or unconventional histology (Methods) by univariate or multivariate analysis accounting for age, sex and tumour grade (Cox proportional-hazards model; *P*>0.05; log-rank test).

Differential expression analysis between both groups revealed a strong downregulation of genes related to adaptive immunity among a subcategory of MPNSTs (Fig. 2a). Specifically, H3K27me3 loss was strongly associated with high tumour cellularity estimates, decreased immune cell infiltration and adaptive immune response activation, and decreased expression of granzymes (*GZMA*, *GZMK* and *GZMH*) and immune checkpoints (*PD-L1* and *HAVCR2;* Fig. 2a-b, Extended Data Figs. 2d-e and 3). Both differential analysis of genome-wide DNA methylation patterns and gene expression revealed a significant activation of genes involved in neural development and morphogenesis pathways in tumours with H3K27me3 loss, which is consistent with the role of PRC2 in neural development^12^ (Extended Data Fig. 2e,g). Together, these results indicate that MPNSTs arising among individuals with NF1 stratify into two clinically and biologically distinct groups characterised by differential levels of immune cell infiltration.

**Figure 2.**
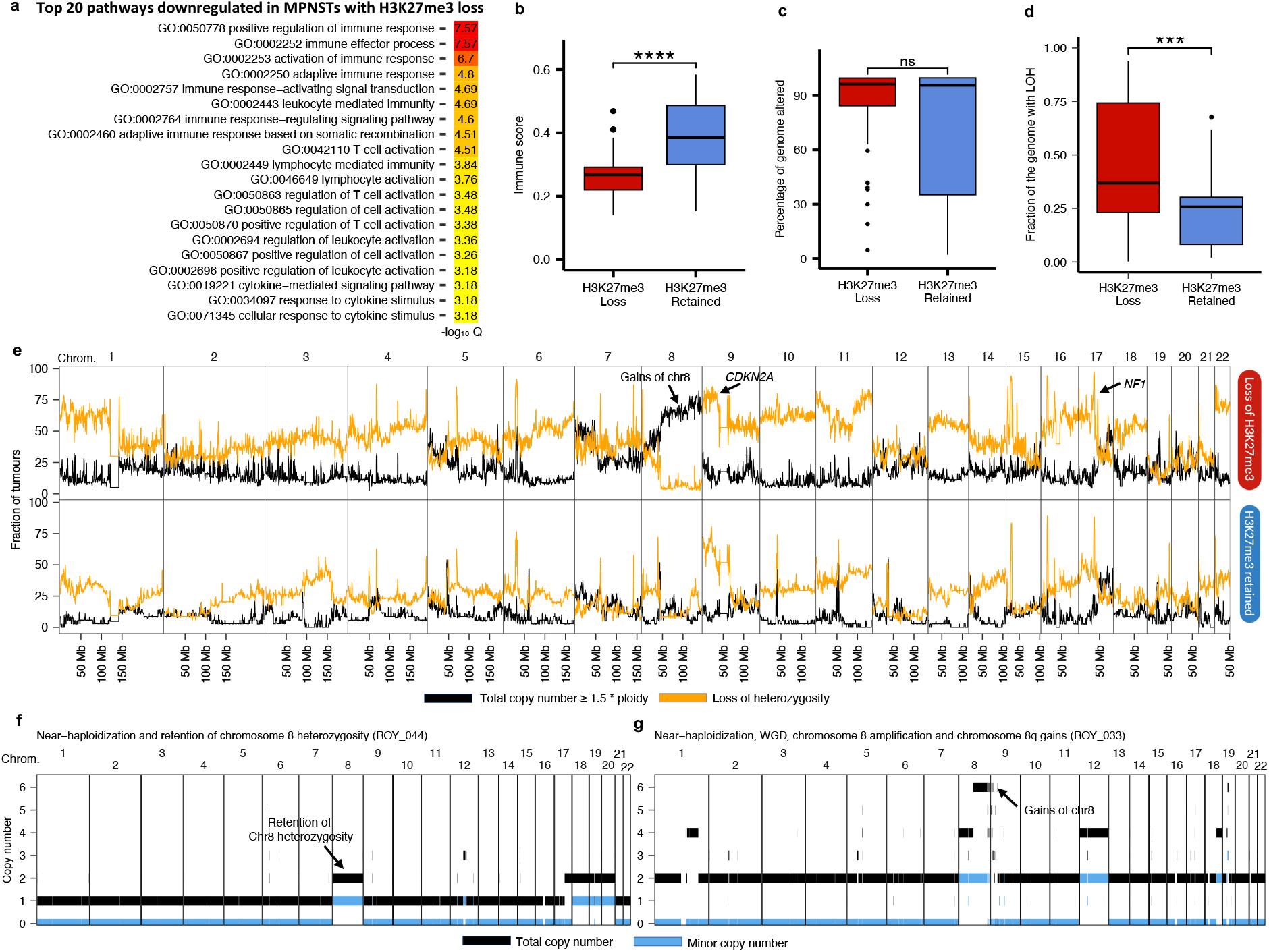
MPNSTs stratify into two clinically and biologically distinct groups according to H3K27me3 status. **a,** Pathways related to immune activation are downregulated in tumours with loss of H3K27me3 estimated. **b**, Immune score analysis reveals decreased immune infiltration according to H3K27me3 status. **c**, Tumours with retention and loss of H3K27me3 are characterised by high levels of somatic copy number alterations. **d**, Tumours with loss of H3K27me3 show significantly higher levels of genome-wide LOH as compared to tumours with retention of H3K27me3. **e**, Genome-wide analysis of the fraction of tumours showing amplification (black) and LOH (orange) based on H3K27me3 status. Representative copy number profiles for tumours with loss of H3K27me3 showing near-haploidization (**f**) followed by whole-genome doubling and chromosome 8q amplification (**g**). Box plots in **b-d** show median, first and third quartiles (boxes), and the whiskers encompass observations within a distance of 1.5× the interquartile range from the first and third quartiles. Amp., amplification; CN, copy number signature; chrom., chromosome; LOH, loss of heterozygosity; WGD, whole genome doubling.

### MPNST evolution is defined by recurrent patterns of copy number alterations

Next, we compared the patterns of MPNST copy number profiles to H3K27me3 status. We found focal deletions of the *CDKN2A* and *NF1* loci and a high burden of SCNAs irrespective of H3K27me3 status. However, both sporadic and NF1-related MPNSTs with loss of H3K27me3 showed a strong enrichment for chromosome 8 amplification and variable levels of genome-wide LOH (Fig. 2c-e, Extended Data Fig. 4, Supplementary Fig. 9 and Methods). LOH of selected chromosomes, including 1p, 10, 11, 16, 17 and 22, was frequently detected in MPNSTs with loss of H3K27me3 (Fig 2e). In one extreme H3K27me3 loss case (Fig. 2f), we observed near-haploidization characterised by genome-wide LOH excluding a few chromosomes with retention of heterozygosity, including chromosome 8, and a ploidy of 1.3. In most other cases, near-haploidization was followed by WGD (Fig. 2g and Supplementary Fig. 9). Copy number signature analysis^27^ identified 18 distinct signatures (Extended Data Fig. 5a) and confirmed an enrichment of signatures related to LOH in MPNSTs with loss of H3K27me3 (Fig. 2e and Extended Data Fig. 5b). In contrast, tumours with retained H3K27me3 were enriched in signatures associated with a diploid genome and genomic instability (*P*=0.029, two-sided Mann-Whitney test; Extended Data Fig. 5b). Together, these results indicate that distinct copy number patterns are associated with H3K27me3 status.

Our analysis of MPNST copy number profiles offers a basis for comparison to other types of cancer. To this aim, we next compared MPNSTs to diverse cancer types from TCGA at the copy number level. Gains in chromosome 8q have also been detected in a number of other solid tumours^28–30^. Numerous cancer-related genes are located on chromosome 8q and have been implicated in cancer progression, including genes such as *C-MYC, RAD21, UBR5*, and *HEY1*^28,31–33^. Of the >700 protein-coding genes on chromosome 8, 199 genes are differentially expressed in the tumours with H3K27me3 loss, including *RAD21,* which has been linked with the mitigation of replication stress caused by the oncogenic *EWS-FLI1* fusion in Ewing sarcomas with chromosome 8q amplification^28^*, UBR5*, and *HEY1* (Supplementary Fig. 10, Supplementary Table 2 and Methods). Near-haploidization has been observed across diverse cancer types, including undifferentiated soft tissue sarcomas, adrenocortical carcinoma, gliomas, chondrosarcomas, and acute lymphoblastic leukemia, and is often linked with a poorer prognosis^25,34–38^. In pan-cancer analysis, MPNSTs with loss of H3K27me3 show the highest levels of genome-wide LOH, including near-haploidization (Extended Data Fig. 6). However, concomitant retention of chromosome 8 heterozygosity and genome-wide LOH as observed in MPNSTs with loss of H3K27me3 (Fig. 2a) has not been linked with a distinct biological process previously. This new copy number configuration provides a potential new avenue to investigate the evolutionary constraints leading to genome-wide LOH^39^.

### Copy number profiles are detected in cell-free DNA and predict patient outcome

Given the strong association between distinct copy number profiles with H3K27me3 status and immune infiltration, we evaluated the clinical relevance of SCNA patterns to predict overall survival in comparison to the predictive ability based on clinical data and H3K27me3 status (Methods). For this analysis, we focused on high-grade MPNSTs arising in NF1 individuals to increase statistical power. Among the set of chromosome arms showing recurrent copy gain, copy loss, or loss of heterozygosity (LOH), LOH of chromosome 5q was the most predictive feature of poor prognosis (Fig. 3a-b). Other copy number aberrations predictive of poor prognosis are LOH of chromosomes 11q, 7p and 22q, and amplification of chromosome 2q and 9q (*P*<0.05; log-rank test). There appeared to be a trend toward worse prognosis among tumours showing amplification of chromosome 8q, although not reaching statistical significance (Fig. 3c). Next, we investigated whether copy number patterns associated with H3K27me3 loss and predictive of survival can be detected in cell-free DNA (cfDNA) collected from a separate cohort of patients with NF1-related MPNST at different time points and analysed by ultra-low-pass whole-genome sequencing^40^ (Methods). Overall, we detected in cfDNA copy number aberrations recurrently found in MPNSTs using WGS data (Fig. 2e), including loss of chromosomes 1p, 5q, 4q, and 22, and chromosome 8 amplification (Fig. 3d, Supplementary Figs 11-13). These results thus suggest that copy number profiles detected in cfDNA could serve as a non-invasive surrogate for H3K27me3 status and prognostication, which warrants further investigation in larger cohorts.

**Figure 3.**
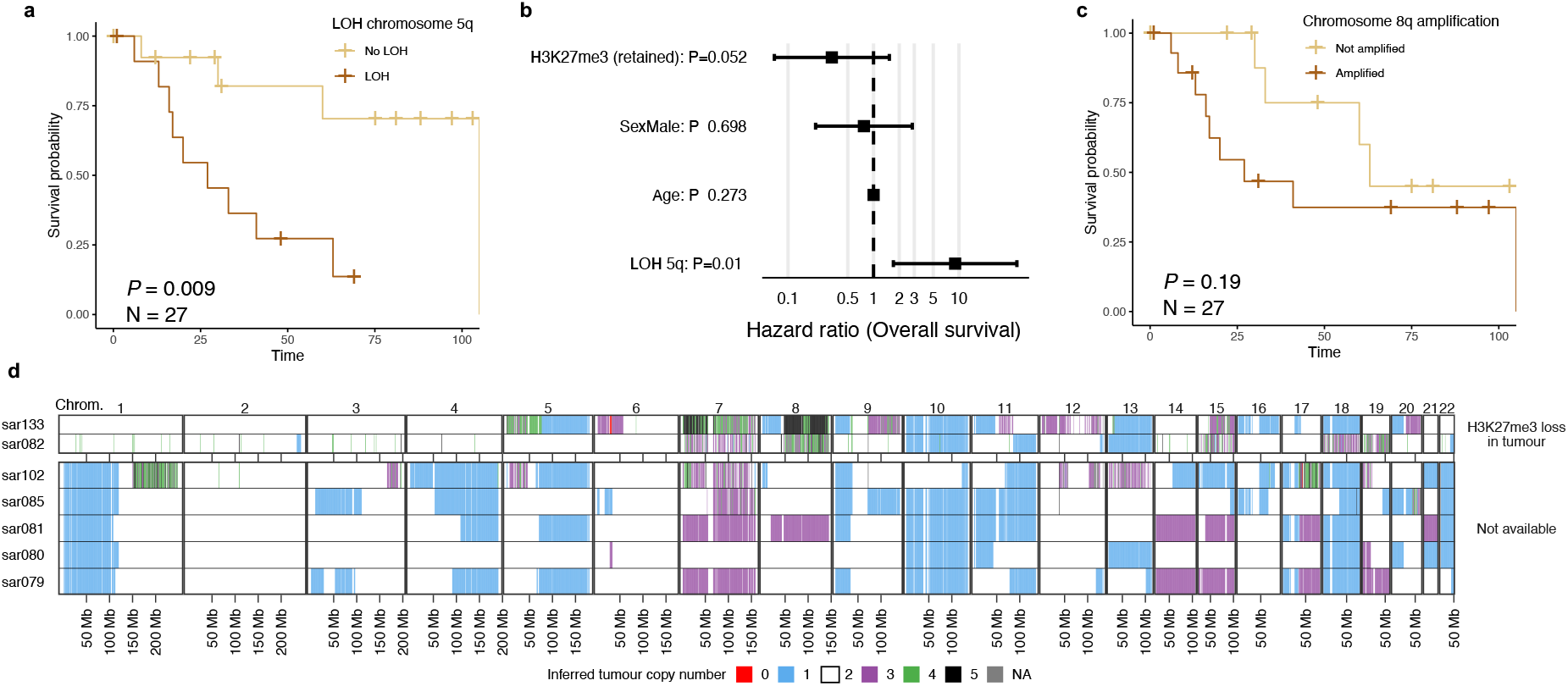
Recurrent copy number aberrations predict patient outcome and are detected in cell-free DNA from NF1 patients. **a**, Kaplan Meier plot showing the overall survival for NF1 individuals with high-grade MPNSTs stratified according to chromosome 5q LOH. **b**, Hazard ratios with 95% confidence intervals and *P* values computed using Cox proportional hazards regression. **c**, Kaplan Meier plot showing the overall survival for NF1 individuals with high-grade MPNSTs based on the amplification of chromosome 8q. **d**, Copy number profiles estimated using ultra-low-pass whole-genome sequencing of cell-free DNA from patients with MPNST.

### Timing of mutations in MPNST evolution

We next sought to determine the relative timing of the genomic alterations that occur during MPNST development across time and space. To this aim, we first investigated which alterations are present in low-grade MPNSTs and MPNST precursor lesions (either benign neurofibromas or transitional lesions referred to as atypical neurofibromatous neoplasms of uncertain biologic potential; ANNUBPs). We detected biallelic inactivation of *NF1* and *CDKN2A* caused by non-recurrent complex SVs in 2/2 neurofibromas, 3/3 ANNUBPs, and 4/5 low-grade MPNSTs (Extended Data Fig. 7). *EED* was mutated by SVs in one neurofibroma and one low-grade MPNST (Extended Data Fig. 7). Analysis of two anatomically distant tumours from the same individual, a neurofibroma and a high-grade MPNST, revealed *CDKN2A* inactivation in both tumours by independent SVs, suggesting parallel evolution and that *CDKN2A* loss is the second genomic alteration after *NF1* inactivation in MPNST pathogenesis (Fig. 4a-c). Together, these results indicate that somatic *CDKN2A* inactivation is an early event in MPNST evolution, which is necessary but not sufficient for the transformation of neurofibromas into MPNSTs.

**Figure 4.**
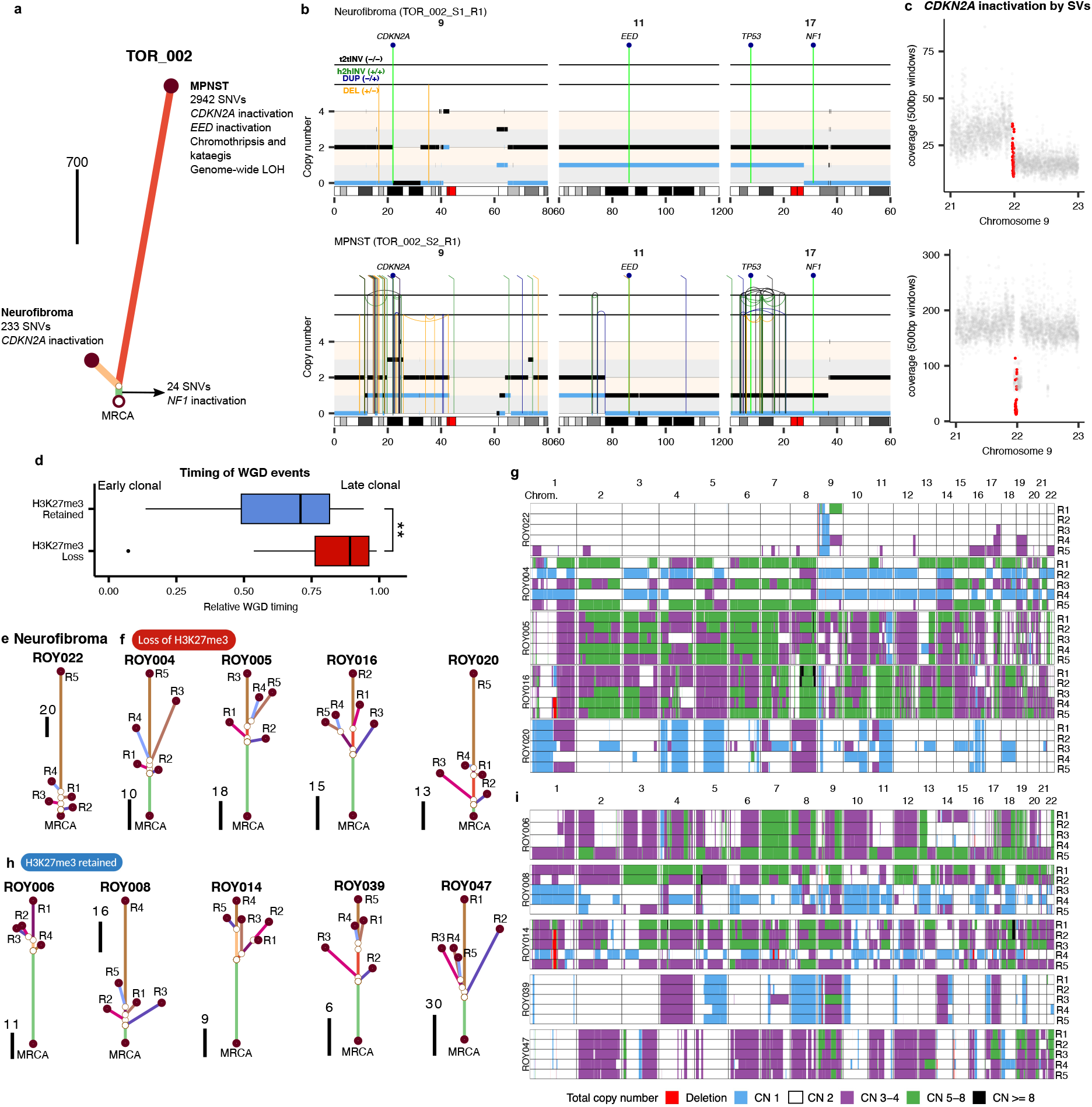
MPNST evolution and intra-tumour heterogeneity. Phylogenetic tree inferred from SNVs detected using whole-genome sequencing data for a neurofibroma and a high-grade MPNST sample from individual TOR_002. Scale bar, 700 SNVs. **b**, Rearrangement profiles of a neurofibroma (top) and a high-grade MPNST (bottom) from patient TOR_002. The genomic position of *CDKN2A, EED, TP53* and *NF1* is shown at the top. Both tumours show biallelic inactivation of *CDKNA* and *NF1* mediated by SVs. Total and minor copy number values are shown in black and blue, respectively. Somatic events common to both tumours (*NF1* deletion and 24 SNVs) and private to each of them are shown. **c**, Sequencing coverage calculated at 500 bp windows at the *CDKN2A* locus for the neurofibroma (top) and the high-grade MPNST (bottom). Dots corresponding to windows mapping to *CDKN2A* are shown in red. **d,** Difference in the timing of whole-genome doubling (WGD) events relative to the clonal expansion between MPNSTs with loss and retention of H3K27me3 (\*\*\**P* < 0.001; two-sided Wilcoxon rank sum test). Phylogenetic trees inferred from SNVs detected using multi-regional sequencing data from fresh-frozen samples for a neurofibroma (**e**), MPNSTs with loss (**f**) and retention (**h**) of H3K27me3. Each region (R) is arbitrarily named with a number. The scale bars indicate the number of SNVs. **g,** Genome-wide total copy number profiles estimated using whole-exome sequencing data for a neurofibroma and MPNSTs with loss of H3K27me3. **i**, Genome-wide total copy number profiles estimated using whole-exome sequencing data for MPNSTs with retention of H3K27me3. MRCA, most recent common ancestor. Box plots in **d** show median, first and third quartiles (boxes), and the whiskers encompass observations within a distance of 1.5× the interquartile range from the first and third quartiles.

Given the high rate of WGD in MPNSTs, we next investigated when WGD occurs during MPNST development. Timing analysis of SCNAs revealed that WGD is consistently a late clonal event in tumours with loss of H3K27me3, and is common in the background of a near-haploid genome (Fig. 4d). In tumours with H3K27me3 loss, WGD is followed by additional gains of chromosome 8q, likely as isochromosome 8q (Methods)^41^. In contrast, in tumours with retention of H3K27me3, WGD occurs earlier in tumour development over a wider period of time (two-sided Mann-Whitney U test, *P*<0.0001; Fig. 4d; and see Supplementary Fig. 14 for an example). These results suggest that WGD events occur at different stages during tumour evolution depending on H3K27me3 status.

To elucidate the patterns of intra-tumour heterogeneity, we performed multi-regional high-depth exome sequencing of five regions from 21 and 15 MPNSTs with loss and retention of H3K27me3, respectively (Fig. 4e-i). Chromosome 8 amplification and WGD were found across multiple spatially distant regions in tumours with loss of H3K27me3 (Extended Data Fig. 8). The late occurrence of WGD events in tumours with H3K27Me3 loss suggests that chromosomal losses, including near-haploidization, may be associated with low fitness that is rescued by the WGD event, which triggers clonal expansion and malignant transformation. Rapid tumour enlargement fosters intra-tumour heterogeneity of mutations and clonal competition, as evidenced by the accumulation of subclonal mutations and variable levels of histopathological heterogeneity in spatially distant regions (Fig. 4h-i, Extended Data Fig. 8 and Supplementary Fig. 15), and might explain the aggressive clinical features of tumours with loss of H3K27me3.

Collectively, our integrative analysis of multi-omic data revealed recurrent patterns of genomic alterations, which allow us to propose clinically relevant mechanistic models of MPNST evolution (Fig. 5). The first steps of MPNST pathogenesis irrespective of H3K27me3 status involve the biallelic inactivation of *CDKN2A,* and in some *TP53* as well, on an *NF1*-deficient genomic background. In MPNSTs with H3K27me3 loss, biallelic inactivation of the PRC2 complex is followed by extensive chromosomal losses, leading to variable levels of genome-wide LOH with retention of chromosome 8 heterozygosity. Late events in tumour evolution include WGD and further amplifications of chromosome 8q, which might be accompanied by additional chromosomal instability (Fig. 5 and Extended Data. Fig. 9). In contrast, tumours with H3K27me3 retention evolve through extensive chromosome instability and chromothripsis, and display more heterogeneous karyotypes (Fig. 5 and Extended Data Fig. 10).

**Figure 5.**
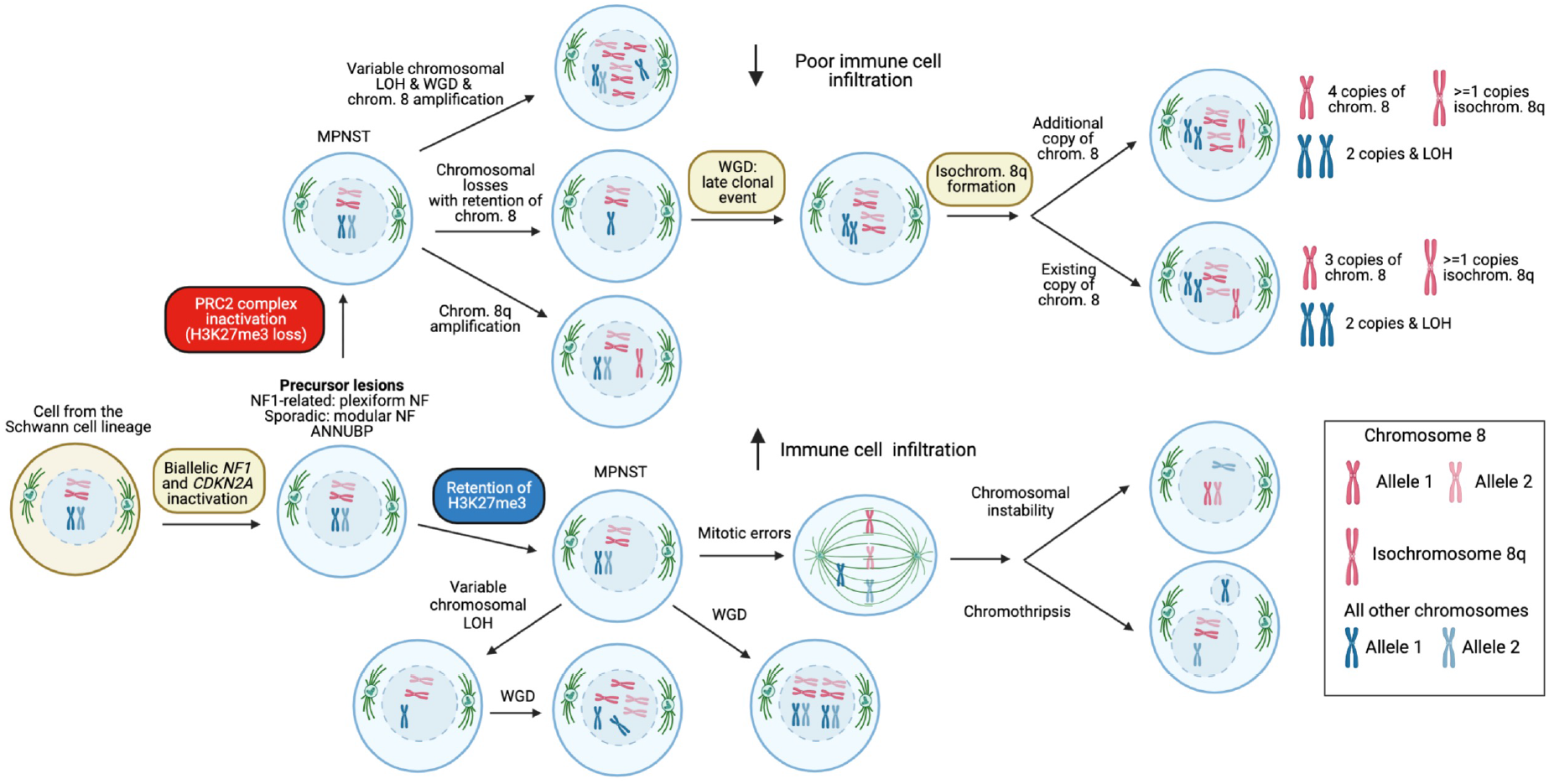
Pathways of MPNST evolution. Schematic representation of the order and timing of events involved in the evolution of MPNSTs. Copy number signatures predominant in specific karyotypic configurations are shown. CN, copy number signature; chrom., chromosome; isochrom., isochromosome; PRC2, polycomb repressive complex 2; WGD, whole genome doubling.

## Discussion

This integrative multi-omic analysis of NF1 and sporadic MPNSTs has expanded the understanding of MPNST development. The international collaboration permitted collection and comprehensive pathologic characterization of a large number of these rare tumours with paired normal samples, thus providing a resource of a large number of these rare tumours with paired normal samples, along with comprehensive pathologic characterization. Multi-omic analysis, coupled with multi-regional deep exome sequencing, revealed distinct genomic evolutionary pathways underlying MPNST pathogenesis that permit subclassification of MPNSTs in a way that correlates with prognosis, and ultimately may lead to individualised treatment approaches. While the focus has been to improve understanding of MPNST development, either in the setting of NF1 or as a sporadic tumour, this expanded MPNST genomic landscape also contributes to the general knowledgebase of other common cancers that share genomic features with MPNST, including loss of *NF1*.

First, we have delineated the order of genomic events leading to this aggressive form of cancer, which has revealed windows of opportunity for intervention through correlation of molecular signatures with tumour status. Our WGS analysis revealed that inactivation of *CDKN2A* is caused by non-recurrent SVs prior to transition from benign to malignant tumours, suggesting that these are random events that increase the fitness of mutant cells. Other loci may contribute to tumour development in a subset of tumours, which may also represent therapeutic targets that could be exploited at an early stage of tumour development. For example, 25% of MPNST cases showed loss of *MTAP*, a gene adjacent to *CDKN2A*. Loss of *MTAP* leads to enhanced dependence on the arginine methyltransferase *PRMT5*^42^, for which there now exist drugs that target this enzyme^43^.

Second, analysis of WGS data revealed that inactivation of the PRC2 complex, which follows inactivation of *CDKN2A*, is mediated by non-recurrent SVs. PRC2 loss is not necessary for MPNST development, as it occurs in only 45% of cases; however, identification of PRC2 loss through genomic analysis of the tumour can provide clinically meaningful information. PRC2 loss correlates with H3K27me3 loss, which in turn correlates with poor prognosis among people with MPNST in the setting of NF1. Thus, genomic analysis of tumour DNA facilitates tumour subclassification into categories predictive of clinical behaviour. Transcriptomic data revealed that tumours with retention of H3K27me3 also show upregulation for markers of immune infiltration and activation of the adaptive immune system, suggesting that this subset of tumours may be more responsive to immunotherapy. Our results suggest that further research into the possibility of incorporating immunotherapy into clinical trials for a subset of MPNST patients is warranted.

Third, we identified a previously unrecognised complex pattern of divergent tumour evolution on the basis of PRC2 loss. In tumours with H3K27me3 loss, biallelic inactivation of the PRC2 complex is followed by chromosomal losses, ranging from loss of several chromosomes to near-haploidization, leading to variable levels of genome-wide LOH with retention of chromosome 8 heterozygosity. Late clonal events in tumour evolution include WGD and further amplifications of chromosome 8q. While chromosome 8 amplification has been found in diverse cancer types, such as Ewing sarcoma, concomitant amplification of chromosome 8 and extensive LOH represents a new copy number configuration specific to MPNSTs with loss of H3K27me3. Tumours with H3K27me3 retention instead evolve through extensive chromosome instability and chromothripsis, and display more heterogeneous karyotypes. Further understanding the transcriptional changes that occur with these copy number aberrations, and comparison of these patterns to other forms of cancer demonstrating similar copy number profiles, may provide insights into novel therapeutic vulnerabilities.

In summary, our results provide the foundation for an MPNST clinical care model that takes into account the genomic architecture of the tumour to provide a more accurate diagnosis, prognosis, and treatment approach. At the level of diagnosis, our comparison of pathologic classification to genomic data clearly demonstrates an inability to identify subsets of tumours based solely on pathology characterization. Our analysis revealed specific genomic patterns of MPNST evolution which enabled classification of tumour subtypes and are more predictive of prognosis than MPNST classification based on clinical or pathologic data alone. Along with more recent data showing the detection of cell-free tumour DNA in plasma from patients with MPNST^40^, the results provide new opportunities for tumour surveillance among patients known to be at increased risk for MPNST development, namely patients with NF1, and in particular those with NF1 caused by a microdeletion^44^. Furthermore, genomic analysis in the clinical setting could benefit all patients with an MPNST through more accurate diagnosis and categorization of these tumours prior to the initiation of either surgical or medical therapy, facilitating a more accurate prognosis and offering an opportunity to assign patients to more specific treatment protocols.

## Methods

### Human Subjects

The Genomics of MPNST (GeM) Consortium is comprised of investigators from academic centres and hospitals (“member sites”) with eligible subjects and existing tissue banking protocols with consent for: biospecimen and clinical data collection; genetic testing, including but not limited to whole genome/exome sequencing; sharing of specimens/data with outside institutions, researchers, etc. The GeM coordinating centre at Boston Children’s Hospital (BCH) coordinated sharing of retrospectively-collected specimens from existing tumour banks and pathology archives at member sites to perform molecular characterization of MPNST and related tumours. The Dana Farber Cancer Institute IRB determined that this research met the criteria for exemption from IRB review and was categorised as Secondary Use research.

All specimens and data were stripped of Protected Health Information (PHI) using the Safe Harbor Method and coded. The prefix of each tumour ID included in the final cohort in Fig. 1 (n=90) is an abbreviation of the member site which contributed the required specimens for analysis: Royal National Orthopaedic Hospital at University College, London (“ROY”, n=43), Moffitt Cancer Center, Tampa (“MOF”, n=12), Mount Sinai Hospital, Toronto (“TOR”, n=9), Massachusetts General Hospital, Boston (“MGH”, n=8), Washington University School of Medicine, St. Louis (“WUS”, n=6), Nagoya University, Nagoya (“NAG”, n=4), Boston Children’s Hospital/Dana Farber Cancer Institute, Boston (“BCH”, n=3), New York University Langone Medical Center, New York (“”NYU”, n=3), Huntsman Cancer Institute at University of Utah, Salt Lake City (“HCI”, n=1), Lifespan Laboratories, Rhode Island Hospital, Providence (“LIF”, n=1).

The coordinating centre at BCH received coded material from the following types of biospecimens: MPNSTs (both sporadic and NF1-related); precursor and other lesions, such as benign, atypical, and plexiform neurofibroma; metastatic and local recurrence neurofibroma; related specimens such as peripheral blood or tissue for germline comparison, and normal nerve where available. Subjects with tumours that were unrelated to MPNST or neurofibroma and subjects with specimens of insufficient quality and/or quality for pathology interpretation were excluded from this study.

Comprehensive clinical and pathology report data for each participant was collected by the international MPNST Registry at Washington University School of Medicine (WUSM). Data include demographic information, disease course, tumour size/anatomical location, histological/immuno-histochemical characteristics, diagnostic imaging, surgical procedures, systemic treatment information, neoadjuvant therapy, toxicity, clinical outcomes and survival.

### Processing of fresh-frozen tumour and paired normal samples

The GeM Consortium’s Standard Operating Procedure (SOP) for the processing of tissue for pathology review and molecular analysis was modelled on the Royal National Orthopaedic Hospital’s SOP for the 100,000 Genomes Project, founded by England’s National Health Service in 2012, as previously described^21^. Fresh-frozen tissue sections (H&E; 5 µm) were assessed to select the most viable areas comprising high quality tumour samples in terms of cellularity, lack of necrosis, and areas with little contaminating non-neoplastic tissue. Fresh-frozen sections of high cellularity (at least 40%) and with less than 20% necrosis were used for nucleic acid isolation. After pathologist assessment, two or four tubes of fresh-frozen tumour tissue scrolls (sectioned at 10um each) were collected depending on whether the tumour was of high cellularity (10,000+ cells) or low cellularity (<4,000 cells), respectively. Each tube (Sarstedt – Biosphere Safeseal Tube 2.0 mL) of fresh-frozen tumour sections for DNA isolation contained 20 curls of tissue in a buffer solution. Two tubes (Sarstedt – Biosphere Safeseal Tube 1.5 mL) with 6 curls each of fresh-frozen scrolls (sectioned at 10µm each) in 1 mL TRIZOL were collected for RNA isolation, regardless of tumour cellularity. Four additional multi-regional samples were taken from a subset of 9 fresh-frozen NF1-related MPNST specimens to assess intra-tumour heterogeneity by performing 500x exome sequencing, whole transcriptome sequencing by RNA-seq, and methylation profiling. Normal tissue samples were confirmed to be free of tumour by a pathologist.

Fresh-frozen tumour and germline DNA samples were extracted and quality was assessed at two pathology hubs, with approximately 50% of GeM samples processed at each site. Boston Children’s Hospital used the Maxwell RSC Tissue DNA Kit for extraction (Promega, Madison, WI) and the Quantus Fluorometer and Picogreen (Promega) for quantification; Royal National Orthopaedic Hospital used QIAamp’s DNA Mini and Blood Maxi kits (Qiagen, Germantown, MD) for extraction (Thermo Fisher Scientific, Waltham, MA) and the Qubit High Sensitivity (HS) and Nanodrop (Thermo Fisher Scientific) for quantification and qualification, respectively. All tumour/normal specimens with >400 ng total DNA (Picogreen, Promega) and a DIN>=6.0 (TapeStation, Agilent) were included in this study. Fresh-frozen tumour and normal nerve samples in TRIzol were stored at –80°C until transfer to the Broad Institute (Cambridge, MA) on dry ice for RNA extraction (AllPrep DNA/RNA/miRNA Universal Kit; Qiagen). RNA quality and insert-size was assessed by Caliper LabchipGXII (PerkinElmer, Billerica, MA) producing a RQS value (equivalent to RIN). RNA quantity was determined by Quant-iT RiboGreen (Thermo Fisher Scientific). Samples which met the minimum required input (250 ng of purified total RNA with RIN >=6.0) were included in this study.

### Processing of FFPE samples

Formalin-fixed, paraffin-embedded (FFPE) tissue blocks were collected for each fresh-frozen tumour which underwent multi-omic profiling and were used for pathology review, immunohistochemistry (MYF-4, H3K7me3, S100, Sox10) and additional molecular characterization (i.e., ploidy assay and multi-regional whole-exome sequencing). FFPE haematoxylin and eosin (H&E) slides from available blocks were reviewed by a pathologist to identify the block most representative (i.e., morphologically similar) to the frozen material which underwent multi-omic profiling. These H&E sections were digitally scanned (40x) at each pathology hub by the Deep Lens Biomedical Imaging Team and uploaded to the VIPER digital pathology platform (Deep Lens, Inc., Columbus, OH). Multi-regional 500x exome sequencing was performed on 5 samples each from 36 FFPE NF1-related MPNST specimens to assess intra-tumour heterogeneity.

### Pathology review

All cases sent for multi-omic profiling underwent expert pathology review in the cloud-based digital imaging platform VIPER hosted by Deep Lens (Columbus, OH). Pathology reports were reviewed for the patient’s NF1 status and lesion’s proximity to a nerve, exposure to neoadjuvant systemic or radiation therapies, and anatomical location. The number of slides available for each case ranged from 1-5 slides, including the following stains: H&E, H3K27me3, MYF-4, S100, and/or Sox10. Additional immunohistochemistry (HMB45 and Melan-A) was performed to exclude melanoma.

The most representative FFPE H&E slide from each case was reviewed by at least one expert pathologist and the following results were recorded in a case review form (CRF): percent tumour necrosis, degree of cytological atypia, tumour cellularity and purity, mitotic count, presence of lymphocytic infiltrate and atypical mitosis, and final tumour diagnosis based on the slide image. Possible diagnoses included: neurofibroma, neurofibroma with atypia, cellular neurofibroma, ANNUBP, low-grade MPNST, conventional high grade MPNST (hyper- and hypo-cellular fascicles with/without geographic necrosis), conventional MPNST with heterologous elements, non-conventional MPNST with associated conventional low-grade areas, non-conventional MPNST, spindle cell/undifferentiated sarcoma, melanoma, and other (e.g., dermatofibrosarcoma protuberans, synovial sarcoma, epithelioid MPNST, spindle cell rhabdomyosarcoma, carcinoma etc.). Additional slides on which IHC was performed from each case were reviewed and results were recorded in CRFs.

Consensus review in real-time via screen sharing and conference calls was conducted for all cases in which there was a discrepant diagnosis between pathologists or diagnostics were difficult based on the available H&E and IHC for each case. A second round of consensus review was conducted for all cases in which there was a discrepancy between tumour diagnosis by consensus review and the results of the methylation assay sarcoma classification. For the purposes of collating tumour diagnosis with multi-omic data, tumour diagnoses were collapsed into five main categories based on germline NF1 status and histological features of the neurofibroma: germline/conventional, germline/non-conventional, sporadic/conventional, sporadic/non-conventional, and equivocal. Equivocal cases were excluded from the genomics analysis unless otherwise stated. Neurofibromas were classified using the criteria outlined in histopathologic evaluation of atypical neurofibromatous tumours and their transformation into MPNST in patients with NF1^45^.

### Whole-genome sequencing

Between 350 and 500ng of tumour DNA were used to prepare DNA sequencing libraries using the TruSeq DNA PCR Free 350bp kit, which were sequenced on the Illumina NovaSeq6000 sequencing machine using S4 flow cells to generate 2 x 151 paired-end reads. Quality control for all steps was performed using a DNA Screentape on the Agilent TapeStation system. Raw sequencing reads were mapped to the GRCh38 build of the human reference genome using BWA-MEM^46^ version 0.7.17-r1188. Aligned reads in BAM format were processed following the Genome Analysis Toolkit (GATK, version 4.1.8.0) Best Practices workflow to remove duplicates and recalibrate base quality scores^47^. A minimum sequencing coverage of 30x and 90x was required for the normal and tumour samples, respectively. NGSCheckMate was utilised using default options to verify that sequencing data from tumour-normal pairs corresponded to the same individual^48^.

### Whole-exome sequencing

Four additional multi-regional cores were taken from the most representative FFPE block for a subset of 10 cases for whole-exome sequencing. DNA extraction from FFPE tissue samples was performed using the Covaris truXTRAC FFPE DNA kit with the E220 Evolution focused ultrasonicator system, and Prepito truXTRAC DNA FFPE kit with the Chemagic Prepito-D. Any excess paraffin was trimmed before sectioning an FFPE tissue block, or after the section had been cut from the FFPE block, to maintain an optimal ratio of 80% tissue to 20% paraffin (or higher). Cores of 1.22 mm diameter were taken; any sections or cores longer than 10 mm in length were cut in half before being loaded into the microtubes. The total mass of FFPE samples processed per extraction was between 2 and 5 mg. The quantity and purity of the DNA was assessed using the Qubit dsDNA Assay kit and Nanodrop, respectively (ThermoFisher). The DNA fragment size of each sample was estimated using an Agilent 2200 TapeStation (Agilent Technologies). Exome libraries were prepared using Roche Kapa HyperPlus kit, captured using the HyperExome kit Roche, and sequenced on Illumina NovaSeq6000, to generate 2 x 151 paired-end reads.

### Tumour DNA ploidy assay

Additional scrolls were cut (50 um) from the most representative FFPE block for all cases for a ploidy assay. Of the 95 frozen cases that qualified for histological assessment by pathology review, 14 had insufficient FFPE-derived tumour DNA qualifications for the ploidy assay. Quantification of DNA content was performed as previously described^25,49^. Briefly, 50 mm FFPE sections were deparaffinized and rehydrated, and cytoplasmic digestion with protease type VIII (Sigma P5380) was utilised to create nuclear suspensions. Nuclear suspensions underwent filtering, cytospinning, DNA hydrolysis (5 M HCl) and staining with Feulgen (Schiff’s fuchsin-sulphite reagent; Sigma S5133). Individual nuclei were classified as “normal”, “lymphocyte”, “plasma”, “fibroblast” and “tumour” using the PWS Classifier software^50^. Nuclei were manually revised to select high confidence members of the “normal”,” lymphocyte” and” tumour” categories. The median integrated optical density (IOD) value of the combined “normal” and “lymphocyte” categories was taken as the IOD level for a diploid cell (IOD_2n_) in each sample. The maximum density value of the IOD of the major tumour subclone was taken as the tumour clonal IOD level (IOD_t_). Tumour ploidy was calculated as 2(IOD_t_/IOD_2n_).

### Detection of pathogenic germline *NF1* alterations

The germline short variant discovery workflow from GATK (version 4.1.8.0)^47^, Strelka2 (version 2.9.2)^51^ and VarDict (version 1.8.2)^52^ were used to detect germline single-nucleotide variants (SNVs) and small insertions and deletions (INDELs) in the whole-genome sequencing data from the matched normal samples. In brief, Strelka2 and VarDict were run using default parameter values. In the case of GATK, intermediate GVCF files were generated for each sample using HaplotypeCaller in GVCF mode. Next, GVCF files for all samples were consolidated into a single GVCF file using the GATK functionality CombineGVCFs using default options. Finally, all samples were jointly genotyped using GenotypeGVCFs and default options. Annovar (version 2018Apr16) was used to annotate all variants^53^. Truncating variants detected by at least one algorithm were considered to be pathogenic, and missense variants were reviewed for literature support. All variants were also manually curated through visual inspection of raw sequencing reads using BamSnap^54^. Deep intronic mutations generating cryptic splice sites were validated at the RNAseq level and through literature review. To detect germline structural variants (SVs) we followed two strategies. First, we called SVs in all matched normal samples using Manta (version 1.6.0)^55^, LUMPY (version 0.2.13)^56^, SvABA (version 1.1.3)^57^, and Delly (version 0.8.3)^57,58^. Breakpoints mapping to the *NF1* locus, including 100Kb upstream and downstream, were manually curated by inspecting the raw sequencing reads. Second, to identify *NF1* microdeletions without SV support, we computed the B-allele frequency profile for chromosome 17 for each sample using the heterozygous SNPs from dbSNP (build 151). In addition, we computed the sequencing coverage at 500bp windows using mosdepth^59^. To call an *NF1* microdeletion the B-allele frequency (BAF) and coverage profiles at the *NF1* locus were required to clearly deviate from 0.5 and decrease by a factor of 2, respectively.

### Detection of somatic single-nucleotide variants and indels

Somatic single-nucleotide variants (SNVs) were detected using Mutect2 (GATK version 4.1.8.0), MuSE (version v1.0rc)^60^, and Strelka2 (version 2.9.2) using default options. Somatic indels were called using Strelka2 (version 2.9.2) and Mutect2 (GATK version 4.1.8.0) using default options. Each algorithm was run independently on each tumour sample using the matched normal sample from the same individual as control. Indel calls were left-aligned using the GATK functionality LeftAlignAndTrimVariants. The calls generated by each algorithm were merged using the Python library mergevcf (https://github.com/ljdursi/mergevcf). To maximise specificity, only mutations called by at least two algorithms were considered for further analysis. Mutations overlapping known polymorphisms in dbSNP (build 151) were only considered for further analysis if the coverage in the matched normal sample was at least 20 reads and none of them supported the mutation. Missense variants predicted to be deleterious by MetaLR and MetaSVM as implemented in Annovar (version 2018Apr16) were considered pathogenic. To identify driver genes, we used the R package dNdScv^61^.

### Mutational signature analysis

Mutational signatures were extracted de novo using non-negative matrix factorization (NMF) as implemented in the R package NMF^62^. In brief, for each rank in the set {2,12} 2,000 factorizations were run. The rank maximising the consensus cophenetic correlation coefficient, and for which the residual sum of squares (RSS) showed a clear inflection point, was selected as the optimal rank^63^. The identified signatures were compared against the COSMIC v3.2 catalogue of signatures and assigned to existing ones using a cosine similarity cut-off of 0.85 using the R package MutationalPatterns^64^. Next, the contribution of the set of extracted signatures to each sample was estimated using the function *fit_to_signature*s from MutationalPatterns. For this analysis, signatures appearing in hypermutated tumours were only considered when analysing the tumours.

### Detection of microsatellite instability

Mutations at microsatellite loci were detected using MSIprofiler (https://github.com/parklab/MSIprofiler) as previously described^65^. Only repeats with at least 10 sequencing reads in both the normal and tumour sample were considered for further analysis. The Kolmogorov-Smirnov test was used to compare the distributions of repeat lengths. The level of significance was set at 0.01 after False Discovery Rate (FDR) correction.

### Copy number analysis

For WGS data, the software packages FACETS (version 0.12.1) and ascatNGS (version 2.5.1) were used to detect somatic copy number alterations and to compute purity and ploidy estimates for all tumour samples^66,67^. Both tools were run using default options. The distribution of cancer cell fraction and mutation copy number values for somatic SNVs computed using the copy number values were manually reviewed in all cases to assess the quality of the ploidy and purity calls. In cases where inconsistencies were identified, such as the lack of clonal SNVs, a different combination of purity and ploidy values was selected to refit the copy number profile. The ploidy and purity values were selected to best match the ploidy ascertained from image cytometry ploidy analysis. Timing analysis of whole-genome doubling events was performed as previously described^25^. For WES data, FACETS (version 0.12.1) was used to compute copy number, ploidy and purity calls for each tumour region.

### Copy number signature analysis

For WGS data, allele counts were generated for SNP positions lifted over to hg38 (1,837,383 SNPs) using alleleCounter (https://github.com/cancerit/alleleCount). For WES data, allele counts were instead generated for all hg38 dbSNP v153 common SNPs (12,140,046 SNPs). LogR and BAF values were calculated for SNPs in all normal samples. SNP positions with a high frequency of undetermined genotype (0.68<BAF<0.9 or 0.1<BAF<0.32) were excluded from analysis. Further, any SNPs closer than 75bp to a neighbouring SNP were excluded from analysis. Lastly, SNPs with a high variance in logR were excluded from analysis (SNP logR variance>2*median SNP logR variance). Following SNP removal and cleaning, normal samples with a high variance in logR were removed from the dataset (samples with >1.5*median sample logR variance). LogR and BAF values were calculated for all tumour samples with a corresponding retained normal sample. Tumour LogR values were corrected for GC content, and copy number profiles were generated using ASCAT^68^, with a segmentation penalty of 70, designating SNPs with 0.2<BAF<0.8 in the corresponding normal as heterozygous. For WES samples, segmentation was jointly performed for all samples originating from a single patient using the function *asmultipcf*. Following copy number calling, copy number profiles were summarised as vectors of counts of segments categorised into 48 copy number classes based on combinations of loss of heterozygosity (LOH) status (homozygous deletions, LOH, heterozygous), total copy number (0, 1, 2, 3-4, 5-8 or 9+) and segment length (0-100kb, 100kb-1Mb, 1Mb-10Mb, 10Mb-40Mb, >40Mb). For homozygous deletions, segment lengths were restricted to 0-{100kb, 100kb-1Mb, >1Mb}. See ^27^ for a full description of the development of these categories.

For WGS data, non-negative matrix factorisation as implemented in sigProfilerExtractor was employed to extract de novo copy number signatures from sequential subsets of the dataset as previously described^69^: all samples (n=81), excluding hypersegmented samples (nsegs<1000; n=79), remaining poorly explained samples (cosine similarity<0.85; n=35), samples with a high proportion of LOH (pLOH>; n=12). In each case, the de novo signatures were decomposed into previously identified pan-cancer copy number signatures^27^ using the function *sigProfilerSingleSample*. For hypersegmented samples, a novel signature was identified, for which the attribution was taken from the all-samples extraction. For the remaining samples, the decomposition that best explained the data (highest cosine similarity) was retained. For WES data, samples were decomposed into previously identified pan-cancer copy number signatures using sigProfilerSingleSample^27^.

### Detection of somatic structural variants

Somatic SVs were detected in whole-genome sequencing data using Manta (version 1.6.0), LUMPY (version 0.2.13), SvABA (version 1.1.3), and Delly (version 0.8.3). Each algorithm was run independently on each tumour using the bulk sequencing data from the same individual as control. The calls generated by each algorithm were merged using the Python library mergevcf allowing 200bp of slop at the breakpoints. Only calls generated by at least two algorithms were kept for further analysis. Intrachromosomal SVs were classified into four groups (DEL: deletion; DUP: duplication; h2hINV; head-to-head inversion; and t2tINV; tail-to-tail inversion) depending on the read orientation at the breakpoints following the notation established by PCAWG^70^. SVs detected in at least two tumour samples from different individuals were discarded, as these are likely germline polymorphisms or artefacts. Finally, SVs with at least one breakpoint mapping to telomeres, centromeres, heterochromatin regions, or to blacklisted regions by the ENCODE project were removed^71^.

### Detection of chromothripsis

Chromothripsis events were detected using ShatterSeek (version 0.7) using recommended cut-off values^72^. We made a chromothripsis call if one of the following sets of criteria were satisfied: (1) at least 6 interleaved intrachromosomal SVs, the copy number for at least 7 contiguous genomic segments oscillates between 2 total copy number states, the fragment joins test, or either the chromosomal enrichment or the exponential distribution of breakpoints test; or (2) at least 3 interleaved intrachromosomal SVs and 4 or more interchromosomal SVs, 7 adjacent segments oscillating between 2 total copy number states and the fragment joins test. An FDR of 0.2 was used as the threshold for statistical significance.

### Rearrangement signatures

Intrachromosomal structural variants were classified into categories according to (1) the SV type as determined from the read orientation at the breakpoints, i.e., deletions, duplications, and inversion (Ih2hINV and t2tINV SVs were grouped together); (2) and the size of the genomic region bridged by the breakpoints: 10-10,000bp, 10-100Kb, 100Kb-1Mb, 1-10Mb, and >10Mb. All translocations were grouped in a single category. In addition, SVs were further stratified based on whether they mapped to an SV cluster. To define SV clusters, we used piecewise constant regression applied on the inter-breakpoint distance values sorted by genomic coordinates. We required at least 10 breakpoints per segment (kmin=10) and used a value of 25 for the gamma parameter, which controls the smoothness of the segmentation^26^. SVs with at least one breakpoint mapping to a segment with an average inter-breakpoint distance smaller than 10% the average inter-breakpoint distance were considered to be involved in a cluster. Therefore, SVs were classified into 32 categories: 30 for intrachromosomal SVs and 2 for translocations, depending on whether at least one of the breakpoints participated in a cluster of SVs as defined above. Rearrangement signatures were extracted using NMF as implemented in the R package NMF^62^. As in the case of SBS, the rank in the set {1..12} maximising the consensus cophenetic correlation coefficient and for which the RSS showed an inflection point that was considered optimal. In addition, the RSS difference obtained for factorizations performed on the original and randomised data using the function *randomise* were compared to ensure that the results obtained were not due to spurious correlations^73^.

### mRNA sequencing

At least 250 ng of purified total RNA with RIN >=6.0 were required. Library preparation was conducted using the Illumina TruSeq Strand Specific Large Insert RNA kit (50M pairs) v1. Thermo Fisher ERCC RNA controls were added prior to Poly(A) selection, providing additional control for variability, including quality of the starting material, level of cellularity, RNA yield, and batch to batch variability. Libraries were sequenced on the Illumina HiSeq4000 sequencing machine at the Broad Institute (Cambridge, MA). Sequencing reads were mapped to the transcriptome using the STAR aligner (version 2.7.4a)^74^. Gene expression counts were generated using HTSeq (v.0.6.1p1)^75^ and normalised to transcripts per kilobase million (TPM). GENCODE v22 was used as the gene annotation reference. Differential expression analysis was performed using DESeq2^76^. Gene fusions were detected using ARRIBA (version 2.1.0)^77^, TopHat-fusion (version 2.1.0)^78^, EricScript (version 0.5.5)^79^, and STAR-Fusion (version 1.10.1)^80^. Only those fusions called by at least 2 algorithms or with WGS support were considered for further analysis. Immune infiltration was estimated using consensusTME using the sarcoma cell-type-specific genes derived from TCGA^81^. Immune gene sets were obtained from previous studies^82–84^.

### Methylation array profiling

Tumour DNA derived from fresh-frozen tumour and paired normal nerve samples was used for whole-genome DNA methylation array profiling using the HumanMethylationEPIC beadchip platform as described previously^85^. In brief, 350ng of DNA were bisulfite converted using the Zymo EZ DNA methylation Gold kit (Zymo Research Corp., Irvine, CA, USA) as per manufacturer’s recommendations. Bisulfite-converted samples were processed and hybridised to the Infinium HumanMethylationEPIC beadchip arrays according to the manufacturer’s recommendations. Raw Methylation EPIC data were uniformly processed using the R package minfi (version 1.34.0)^86^. Probes that did not satisfy a detection *P* value of 0.01 were discarded using the function *detP*. Probes mapping to known polymorphisms were identified using the function *dropLociWithSnps* and were discarded. Functional normalisation after background correction using normal-exponential deconvolution using out-of-band probes (noob), as implemented in the function *preprocessFunnorm*, was applied to normalise methylation intensities^87,88^. The function *getBeta* was used to convert methylation intensities to Beta values, which were used for further analysis. Quality assessment was performed through visual inspection of the distribution of B values for all samples. For methylation-based diagnosis, iDAT files were analysed using the open-source DNA methylation sarcoma classifier as described previously^22^.

### Integrative analysis of multi-omic data

We performed clustering of each data type separately using ConsensusClusterPlus^89^. The top 5,000 most variable genes (normalised counts) and the top 10,000 methylation CpG sites (Beta values) based on median absolute deviation were used for clustering analysis. Clustering was performed using the hierarchical clustering algorithm with 1,000 resamples, Pearson correlation as the distance metric, and a maximum number of 10 clusters. Both the analysis of methylation and RNAseq data revealed two main groups that were consistent across data types. The cluster assignments computed using *ConsensusClusterPlus* were used for Cluster-Of-Clusters Analysis (COCA)^90^. COCA, as implemented in the R package coca^91^, was run on the same data sets used for *ConsensusClusterPlus* clustering and using the *k*-means algorithm as the clustering method.

### Analysis of cell-free DNA sequencing data

Sequencing reads were filtered to keep read pairs corresponding to DNA fragments between 80 and 160bp. Genome-wide sequencing coverage was estimated by counting the number of reads mapping to non-overlapping windows of 1Mb using readCounter^92^. IchorCNA^93^ was used to estimate the tumour fraction in cell-free DNA and genome-wide copy number profiles. Only samples with a tumour fraction of at least 5% were considered for further analysis. IchorCNA was run using default parameter values without accounting for subclonal events.

### Survival analysis

Survival analysis was performed using the Cox proportional hazards model as implemented in the R package survival (version 2.30)^94^. Significance was assessed by the likelihood ratio test using a cut-off for statistical significance of 0.05. We only considered MPNSTs with conventional histology for survival analysis. The proportional hazards assumption was tested using the *cox.zph* function from the R package survival^94^.

### Phylogenetic analysis of exome-sequencing data

We constructed a binary matrix for each tumour with rows indexed by somatic SNVs and columns by tumour regions such that the *i,j* entry in each matrix was set to one if mutation *i* is present in region *j*, and to zero otherwise. To reconstruct phylogenetic trees for FFPE cases we used the parsimony ratchet method (*pratchet*) as implemented in the R package phangorn^95^. The *acctran* method was used to infer branch lengths for all trees. In the case of fresh-frozen samples, we used the maximum parsimony and neighbour joining algorithms as implemented in the R package MesKit^96^ to estimate consensus phylogenetic trees for each tumour using 100 bootstrap resamples.

### Data availability

Raw sequencing data has been deposited at the European Genome-phenome Archive (EGA), which is hosted by the EBI and the CRG, under the dataset accession number EGAD00001008608. Ultra-low-pass whole-genome sequencing data of cell-free samples is available under controlled access data at https://doi.org/10.7303/syn23651229.

### Code availability

The code used to process the raw data is available upon request.

## Acknowledgements

This research was funded by the Neurofibromatosis Research Initiative (NFRI) at Boston Children’s Hospital, made possible by an anonymous donation. We thank S. Cano, J. Da, M. Fournier, N. Gokgoz, J. Hart, J. Johnson, D. Lucente, T. Restrepo, M. Roche and A. Silva for technical assistance in collection and processing of specimens. We thank C. Clinton and A. Ward for their guidance and support. I.C.-C. thanks EMBL for funding. NP is funded through a Cancer Research UK grant (grant no: 18387). NP and AMF are supported by the UCLH Biomedical Research Centre and the CRUK Experimental Cancer Centre. Provision of patients’ samples from the RNOH was made possible through the Royal National Orthopaedic Hospital Research and Development Department, The Rosetrees Trust, Skeletal Cancer Trust, Sarcoma UK and The Bone Cancer Research Trust over the last two decades. CDS is funded through Cancer Research UK and the Neurofibromatosis Research Initiative (NFRI) at Boston Children’s Hospital-GEM consortium. DNA methylation profiling at NYU is in part supported by grants from the Friedberg Charitable Foundation, the Sohn Conference Foundation, and the Making Headway Foundation (to M.S.). We thank the patients and their families for their participation in this study.

## Author Contributions

A.A., M.M.B., B.C.D., A.M.F., J.H., A.C.H., K.J., J.T.J., R.H.K., D.L., D.T.M., Y.N., M.S., D.V. and X.W. provided samples. D.C.B., V.E., A.M.F., J.H., A.C.H., J.T.J., D.L., D.T.M., Y.N., K.P., M.S., R.T.S., J.J.S., D.V. and X.W. provided clinical data. A.A., M.M.B., B.C.D., A.M.F., A.C.H., D.L., K.P., J.S. and M.S. provided technical support in specimen processing. D.C.B., V.E., A.C.H. and K.P. conducted clinical data processing. R.T.S. and J.J.S. provided annotated genomic data. A.C.H., I.C.C., A.F., A.M.F., A.G., C.M., P.J.P., N.P., P.P., J.S., M.S. and C.D.S. conducted genomic data processing. I.C.C., A.A., M.M.B., A.M.F., A.C.H., D.L., P.J.P., D.T.M., N.P., M.S., C.D.S. and X. W. developed the experimental design. A.C.H., I.C.C., A.A., A.C, M.M.B., A.M.F., B.C.D., A.G., A.C.H., P.J.P., D.T.M., N.P., J.S., M.S. and C.D.S. conducted data analysis and interpretation. I.C.C., A.M.F., A.C.H., D.T.M., N.P., K.P., M.S. and C.D.S. drafted the manuscript. A.A.I., D.C.B., M.B., A.C., I.C.C., B.C.D., V.E., A.F., A.M.F., A.G., A.G., J.G., J.H., A.C.H., K.B.J., J.T.J., G.J., R.H.K., D.L., C.M., D.T.M., Y.N., P.J.P., K.P.., N.P., P.P., J.S., M.S., C.D.S., R.T.S., J.J.S., N.J.U., D.V. and X.W. reviewed and approved the manuscript. I.C.C., A.M.F., A.C.H., D.T.M. and N.P. jointly supervised the work.

## Group Authors: The Genomics of MPNST (GeM) Consortium

Davies C, Griffin A, Gusella JF, Jour G, Sakai T, Shao D, and Stemmer-Rachamimov AO

## Supplementary Tables

**Supplementary Table 1.** Clinical information for the GeM cohort.

**Supplementary Table 2.** Differentially expressed genes between MPNSTs with loss or retention of H3K27me3 immunoreactivity.

---

**Extended Data Figure 1.**
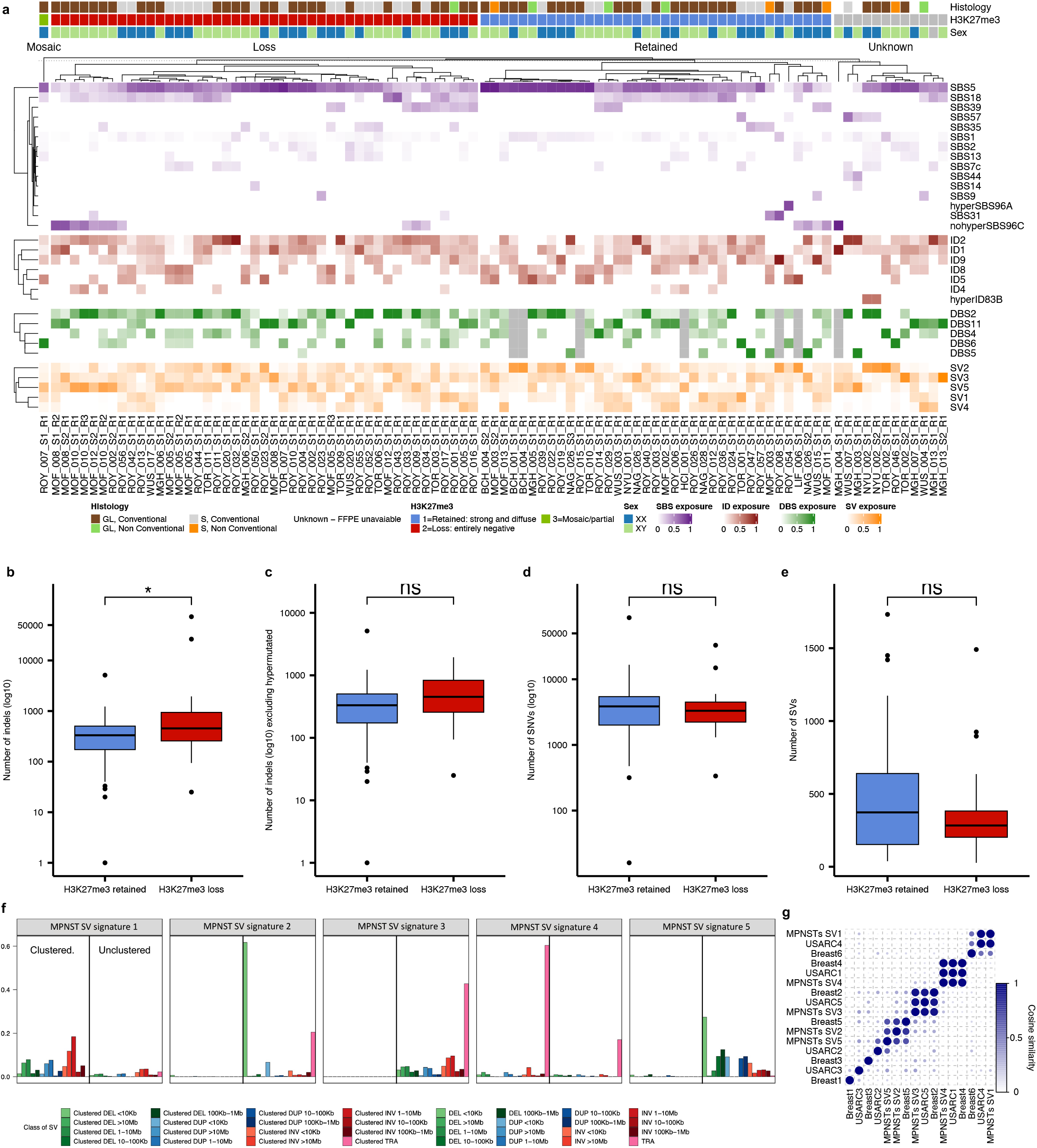
Patterns and rates of somatic mutations in MPNSTs detected using WGS data. **a**, Mutational signatures in MPNSTs. **b,** Genome-wide number of indels. **c,** Genome-wide number of indels excluding hypermutated cases (>20,000 indels). **d** Genome-wide number of SNVs. **e,** Genome-wide number of SVs. **f**, SV signatures detected in MPNSTs. **g**, Comparison of the SV signatures detected in MPNSTs with previously reported SV signatures. Box plots in **b-d** show median, first and third quartiles (boxes), and the whiskers encompass observations within a distance of 1.5× the interquartile range from the first and third quartiles. SBS, single-base substitution signature; ID, indel signature; DBS, doublet base substitution signature; SV, structural variation signature. GL, germline; S, sporadic. ns, not significant, *P* > 0.05; *, *P* < 0.05 (two-sided Wilcoxon rank sum test).

**Extended Data Figure 2.**
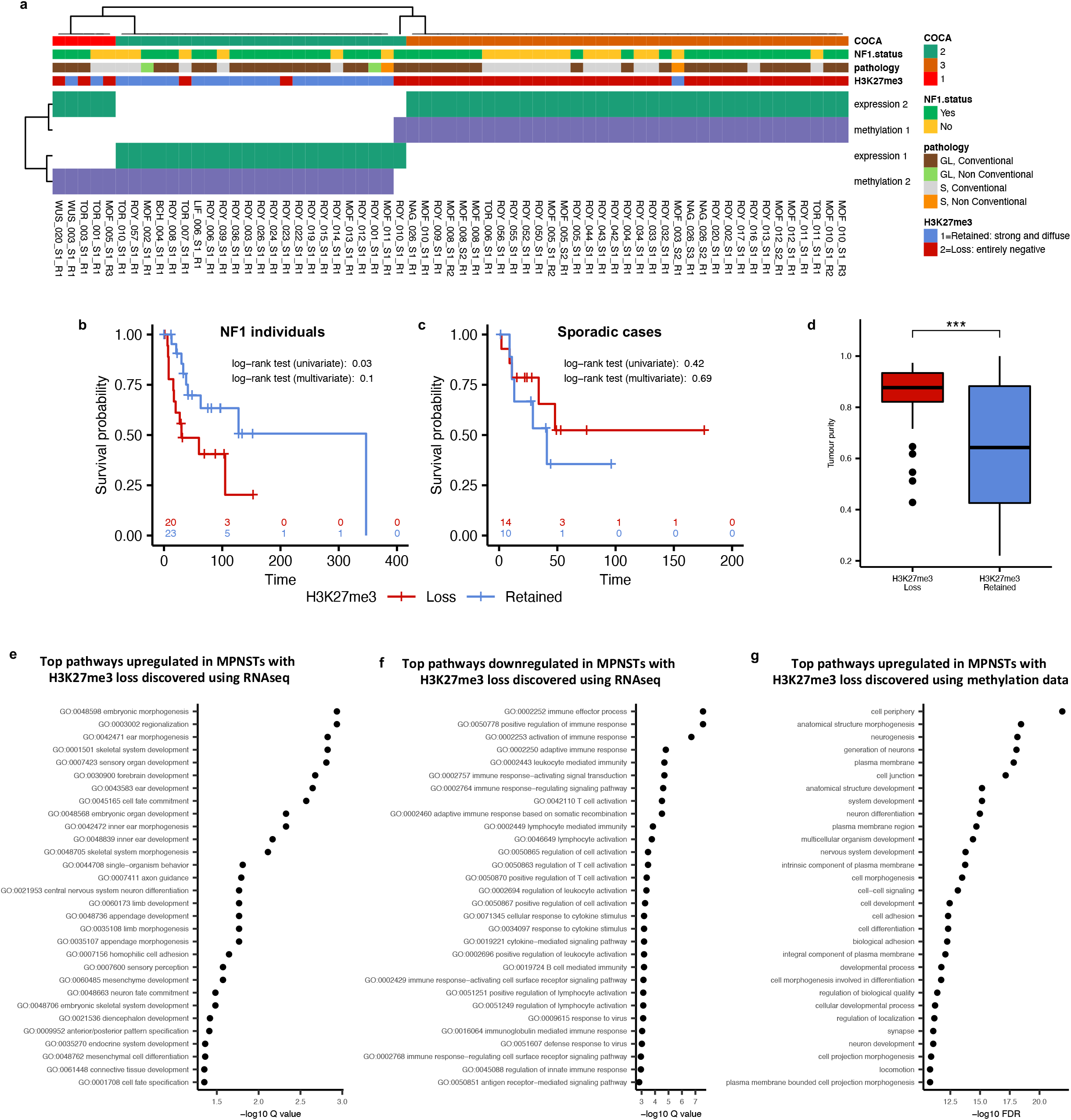
Integrative molecular analysis of MPNSTs. **a,** Cluster-Of-Clusters Analysis (COCA) using methylation and mRNAseq data. **b,** PCA analysis of mRNAseq data. **b**, Kaplan-Meier plots of overall survival of NF1 patients based on H3K27me3 status. **c**, Kaplan-Meier plots of overall survival of patients with sporadic MPNST based on H3K27me3 status. **d,** WGS estimates of tumour purity. Gene Ontology (GO) terms upregulated (**e**) and downregulated (**f**) in MPNSTs with loss of H3K27me based on RNAseq data analysis. **g,** Gene sets enriched in MPNSTs with loss of H3K27me based on the analysis of differentially methylated regions.

**Extended Data Figure 3.**
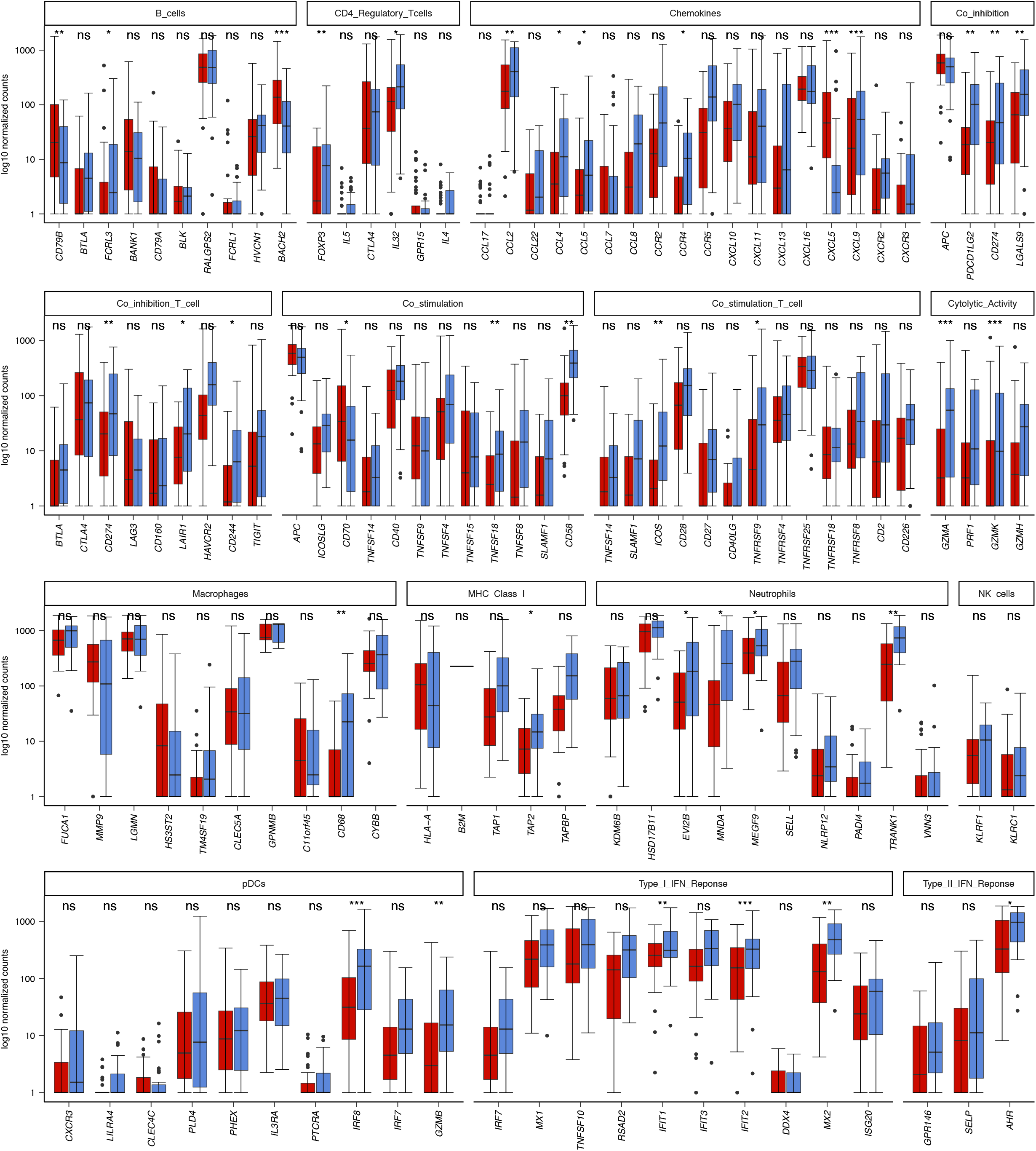
Analysis of the expression of immune signatures in MPNSTs. Box plots show median, first and third quartiles (boxes), and the whiskers encompass observations within a distance of 1.5× the interquartile range from the first and third quartiles. pDCs, plasmacytoid dendritic cells. ns, not significant *P* > 0.05; *, *P* < 0.05; \*\**P* < 0.01; \*\*\**P* < 0.001 (two-sided Wilcoxon rank sum test).

**Extended Data Figure 4.**
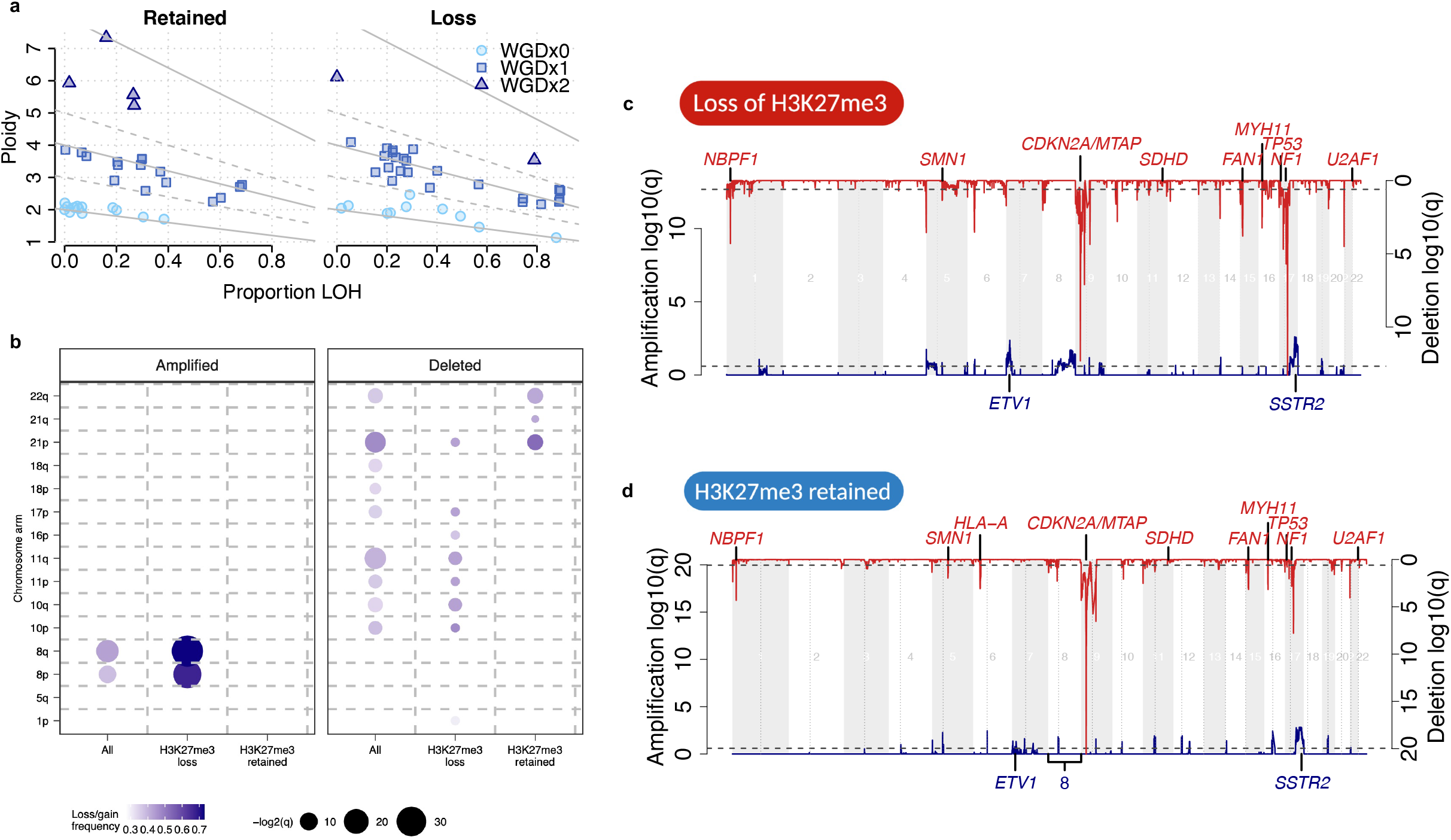
Analysis of recurrent copy number aberrations and whole-genome duplication events. **a,** Relationship between tumour ploidy and the fraction of the genome showing loss of heterozygosity (LOH). **b,** Analysis of chromosome arms recurrently amplified or deleted in MPNSTs. GISTIC analysis of recurrent genome wide copy number amplifications (shown in red) and deletions (shown in blue) in tumours with loss (**c**) or retention (**d**) of H3K27me3, respectively. Genes mapping to segments recurrently amplified or deleted are shown at the top and bottom, respectively.

**Extended Data Figure 5.**
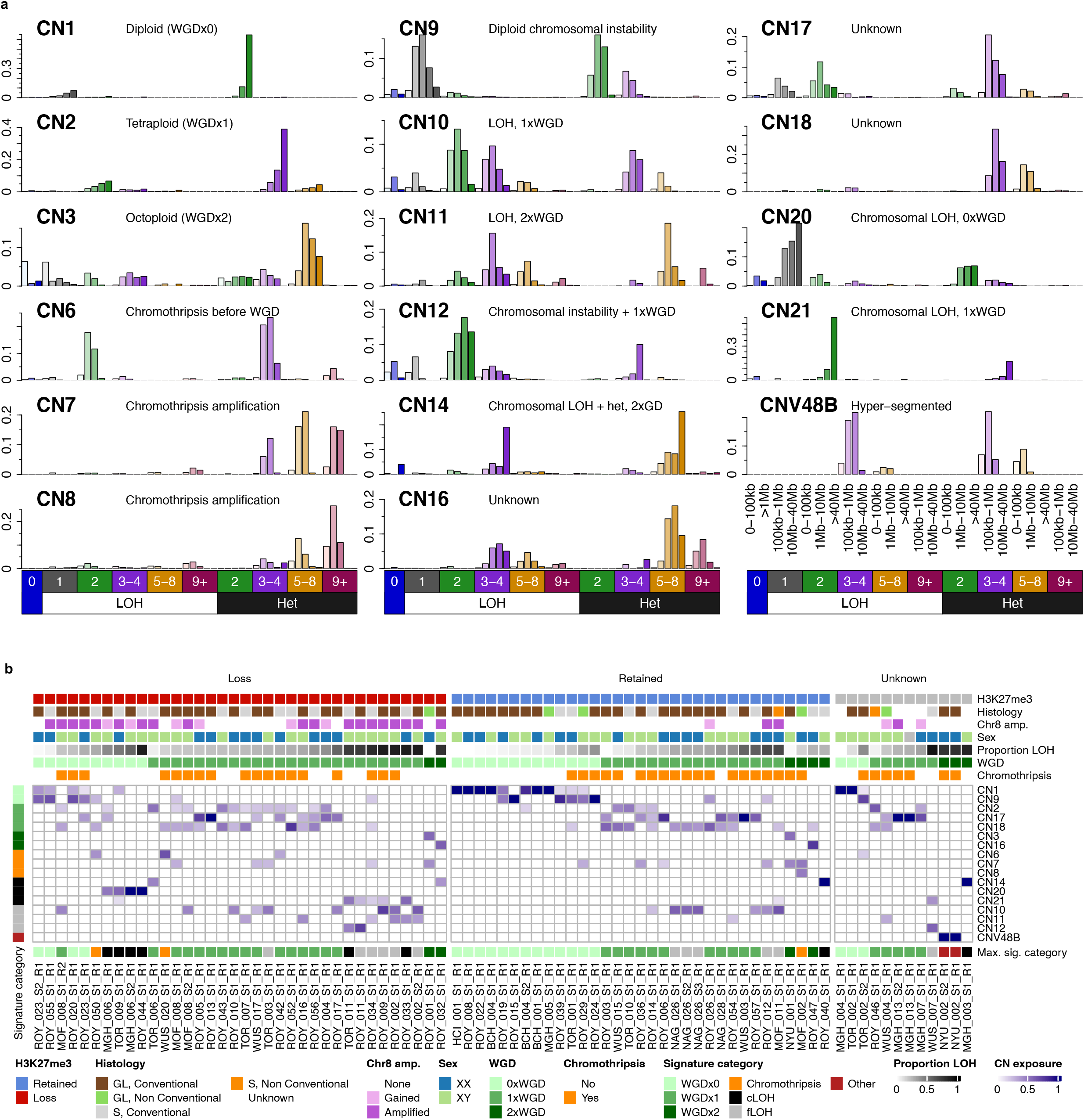
Analysis of copy number signatures. **a,** Copy number signatures detected in MPNSTs using whole-genome sequencing data. CN, copy number signature; LOH, loss of heterozygosity; Het, heterozygous genomic segment; WGD, whole-genome doubling. **b,** The exposure of each copy number signature in each sample is shown. Samples are stratified based on H3K27me3 status. Signature abbreviations. CN1: Diploid (large heterozygous segments with CN=2); CN2: Tetraploid (large heterozygous segments with CN=3-4); CN3: Octoploid (large heterozygous segments with CN=5-8); CN4: Chromothripsis; CN5: Chromothripsis; CN6: Chromothripsis before WGD (LOH segments indicate before WGD); CN7: Chromothripsis associated amplification, after WGD; CN8: Chromothripsis associated amplification, after WGD; CN9: Diploid chromosomal instability; CN10: Extensive LOH, followed by 1x WGD; CN11: Extensive LOH, followed by 2x WGD; CN12: Extensive LOH, followed by 1x WGD, larger segment sizes than CN10; CN13: Early chromosomal LOH, with chromosomal heterozygosity (retained chrs), 1x WGD; CN14: Early chromosomal LOH, with chromosomal heterozygosity (retained chrs), 2x WGD; CN15: Associates with HRD; CN16: Unknown aetiology, 2x WGD, mostly heterozygous; CN17: Unknown aetiology; CN18: Unknown aetiology; CN19: Unknown aetiology; CN20: Early chromosomal LOH, 0x WGD; CN21: Early chromosomal LOH, 1x WGD; CN22: Early chromosomal LOH, 2x WGD; CNV48B: Extreme segmentation.

**Extended Data Figure 6.**
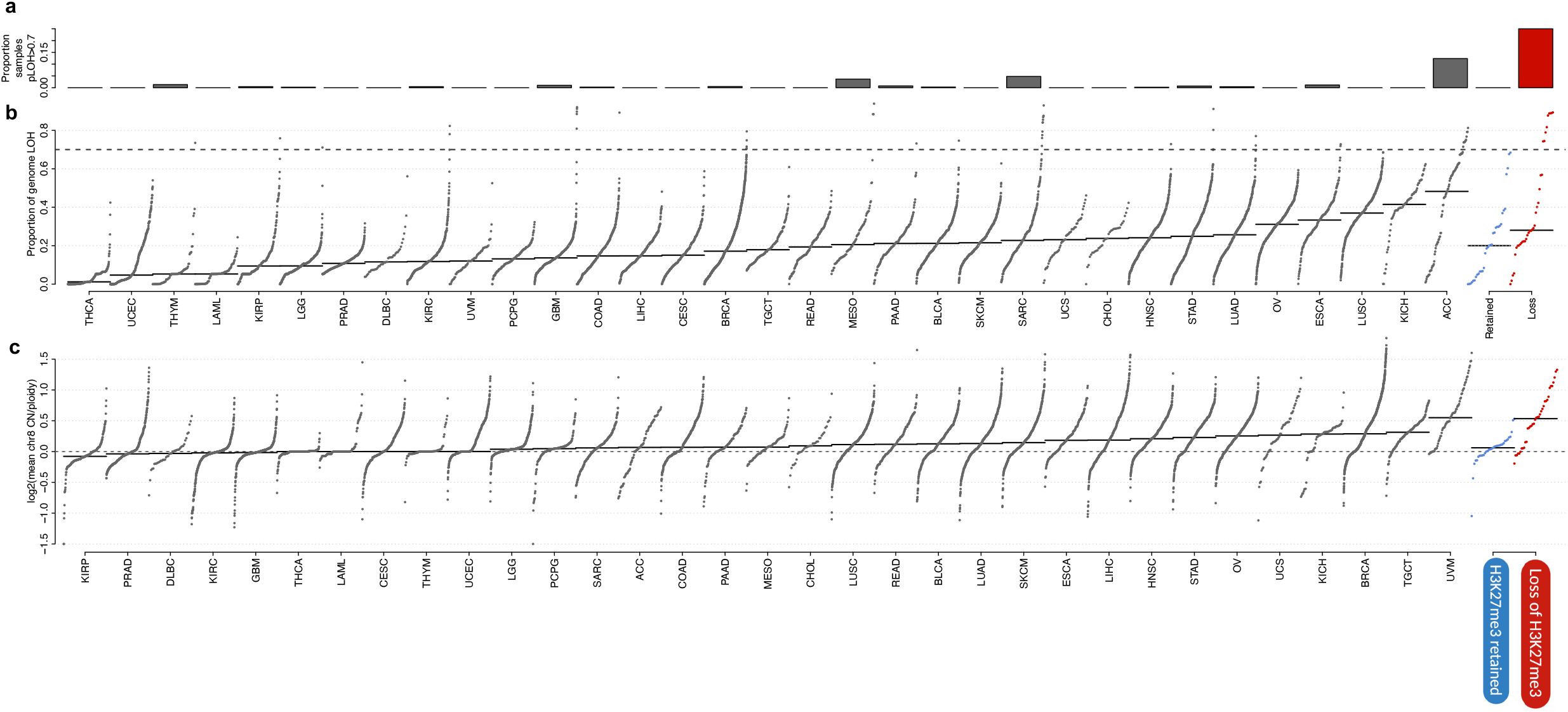
Pan-cancer analysis of genome-wide loss of heterozygosity and chromosome 8 amplification. **a**, Fraction of tumours in TCGA showing LOH in at least 70% of the genome across 33 cancer types. **b**, Fraction of LOH in 9,699 tumours from TCGA. Cancer types are sorted with respect to the 90% percentile (black line). The grey line denotes the median fraction of LOH for each cancer type. **c**, Chromosome 8 copy number in relation to overall ploidy across TCGA. Cancer types are ordered by their median chromosome 8: ploidy ratio (horizontal lines).

**Extended Data Figure 7.**
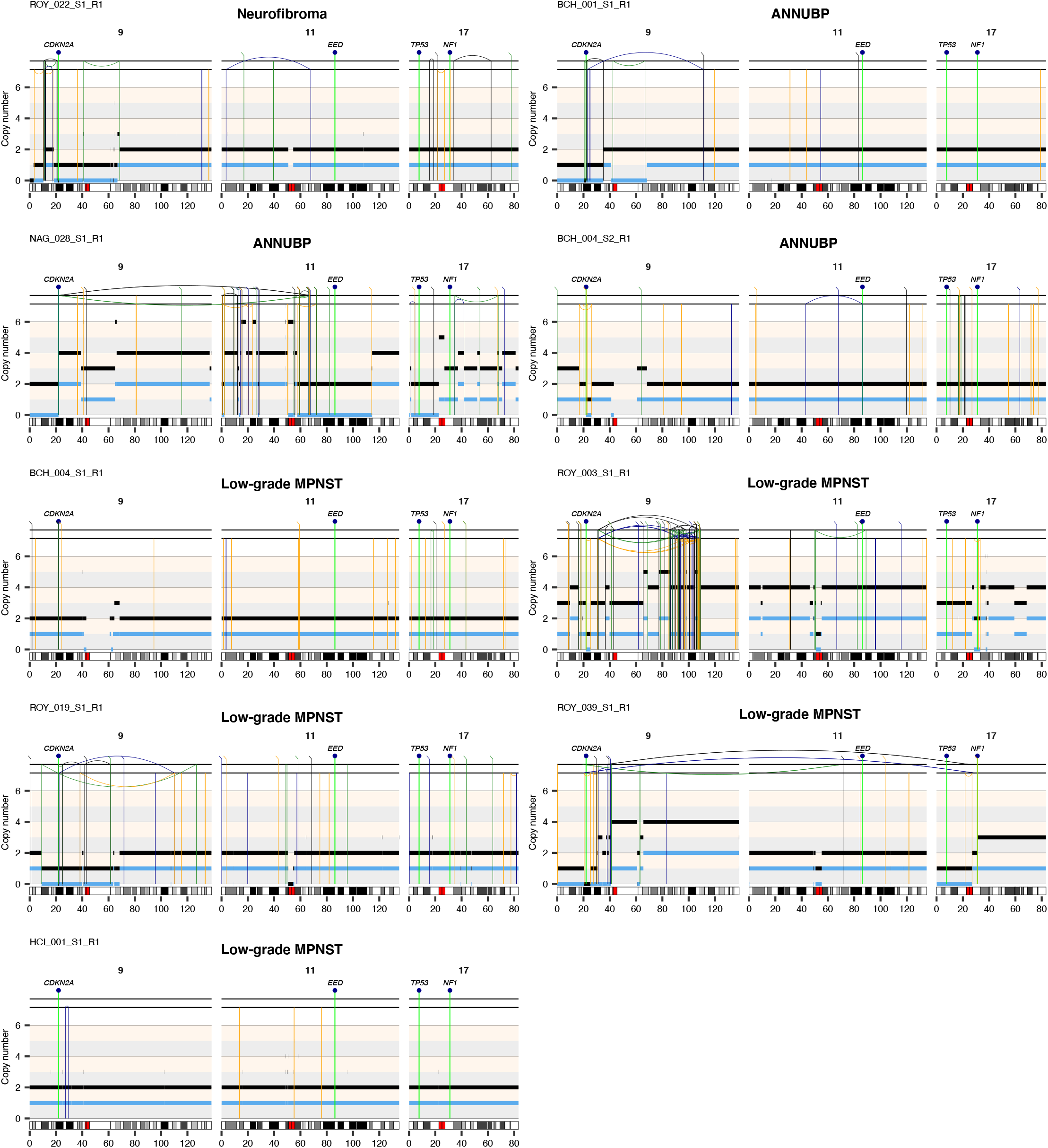
Analysis of rearrangements at the *CDKN2A, EED, TP53* and *NF1* loci. Rearrangement profiles of 1 neurofibroma (ROY_022_S1_R1), 3 ANNUBPs (BCH_001_S1_R1, BCH_004_S2_R1 and NAG_028_S1_R1) and 5 low-grade MPNSTs (BCH_004_S1_R1, ROY_003_S1_R1, ROY_019_S1_R1, ROY_039_S1_R1, HCI_001_S1_R1) in the GeM cohort. The position of *CDKN2A, EED, TP53* and *NF1* is shown at the top of each plot.

**Extended Data Figure 8.**
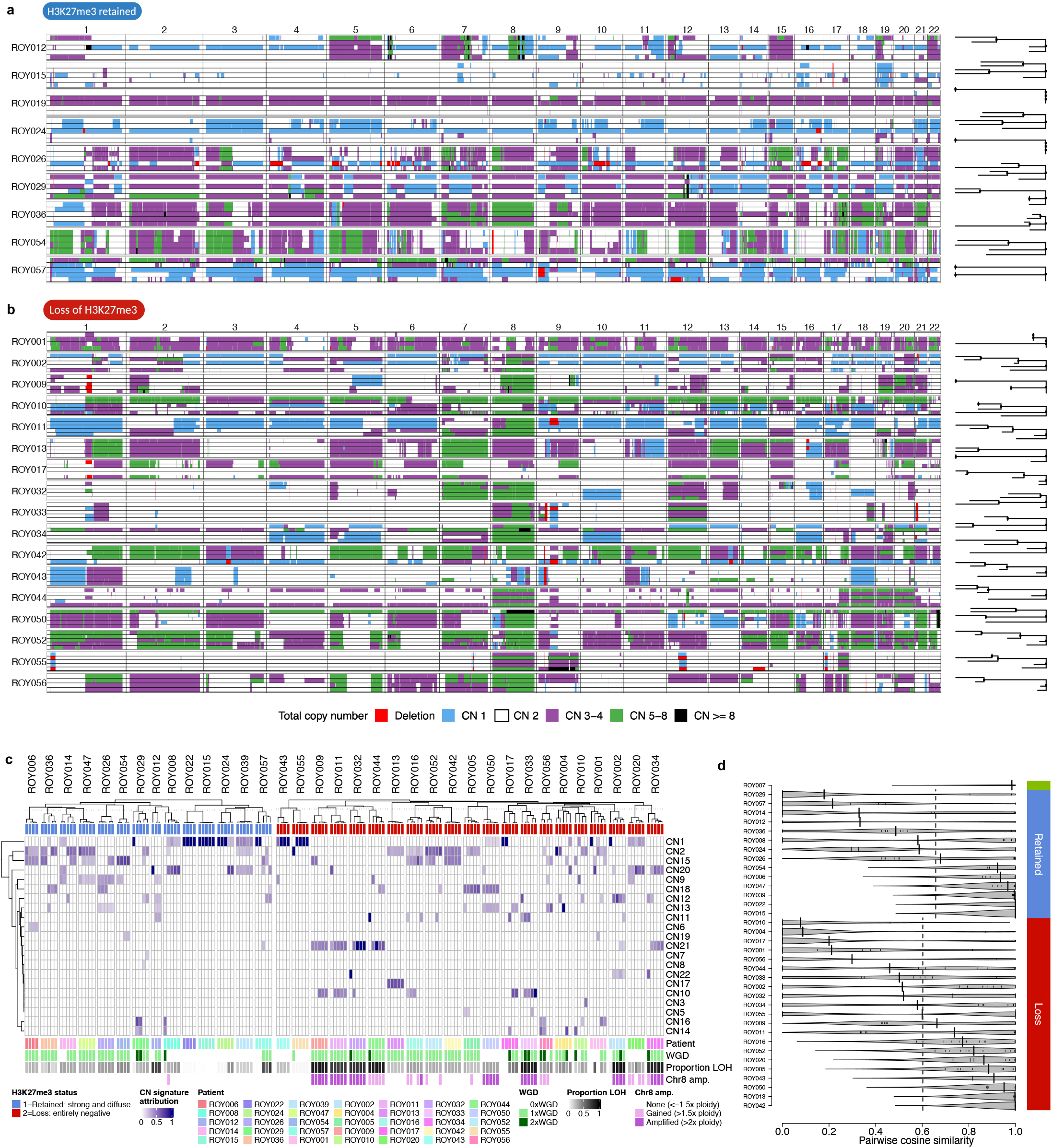
Analysis of the intra-tumour heterogeneity of copy number aberrations in MPNSTs with retention of H3K27me3. Genome-wide total copy number profiles for each region from tumours with retention (**a**) and loss (**b**) of H3K27me3 are shown. The phylogenetic trees generated using SNVs are shown on the right-hand side. **c**, Attribution of copy number signatures in multi-regional exome sequencing data (blue), clustered both within and between patients. Clinical and genetic covariates are shown at the top and bottom of the heatmap. **c**, Pairwise cosine similarity of the copy number signature attributions across regions from the same patient. Individual cosine similarities (thin black lines) and median cosine similarity (thick black lines) are denoted for individual patients. Dashed lines indicate mean pairwise cosine similarities across H3K27me3 loss and retained patients separately.

**Extended Data Figure 9.**
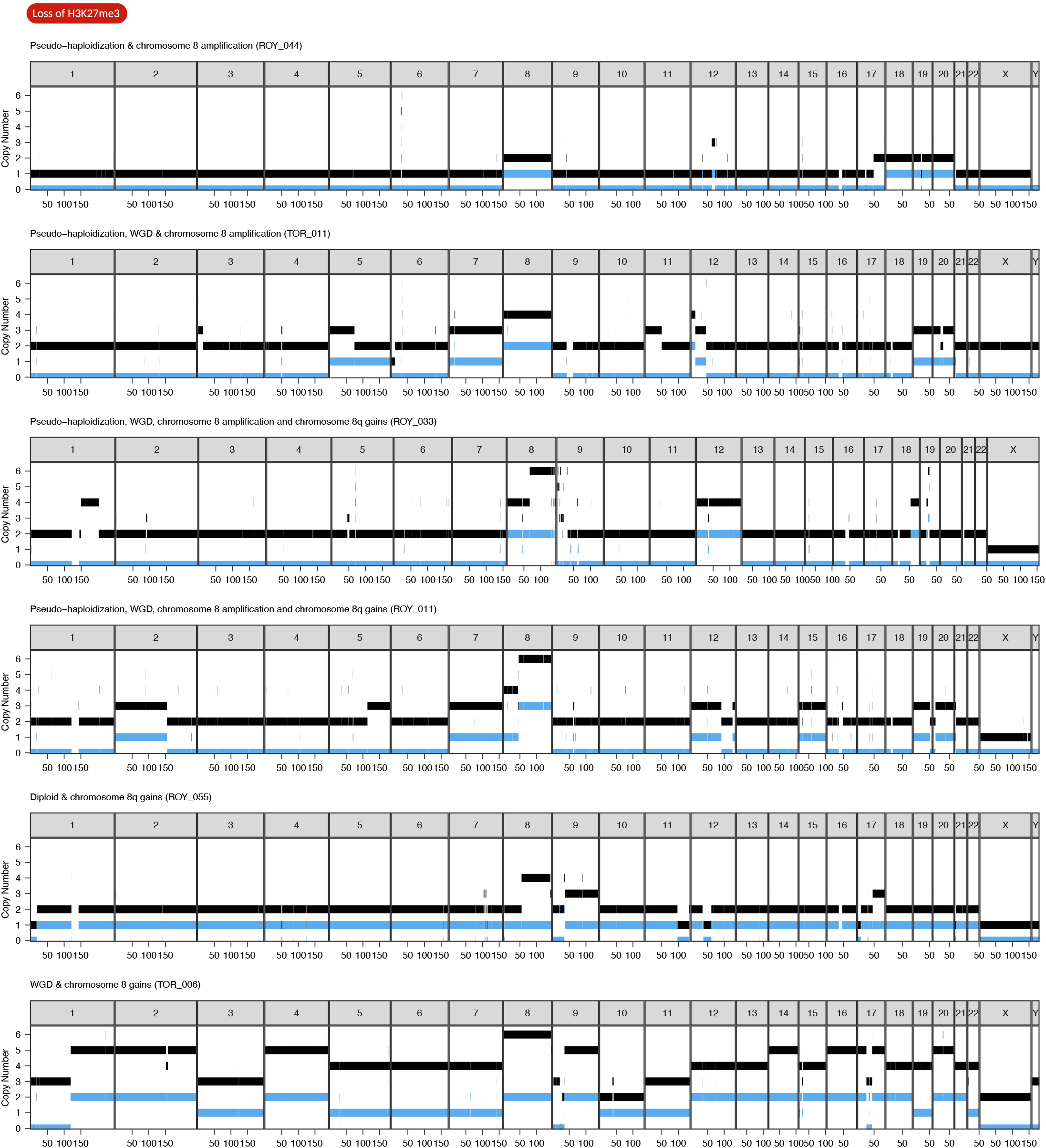
Representative examples of the copy number profiles observed in samples with H3K27me3 loss. Total and minor copy number values are shown in black and blue, respectively.

**Extended Data Figure 10.**
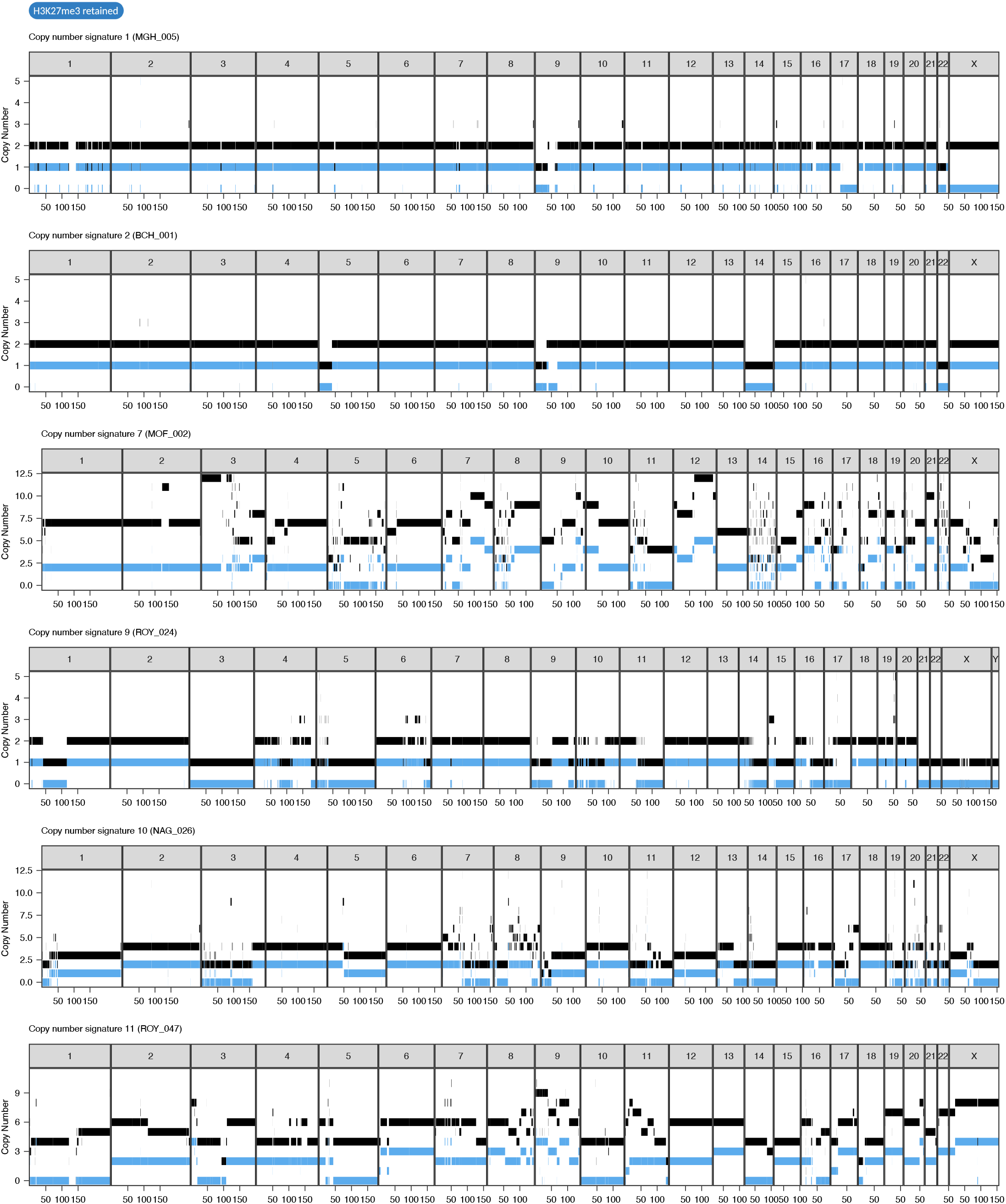
Representative examples of the copy number profiles observed in samples with H3K27me3 retained. Total and minor copy number values are shown in black and blue, respectively.

**Supplementary Figure 1.**
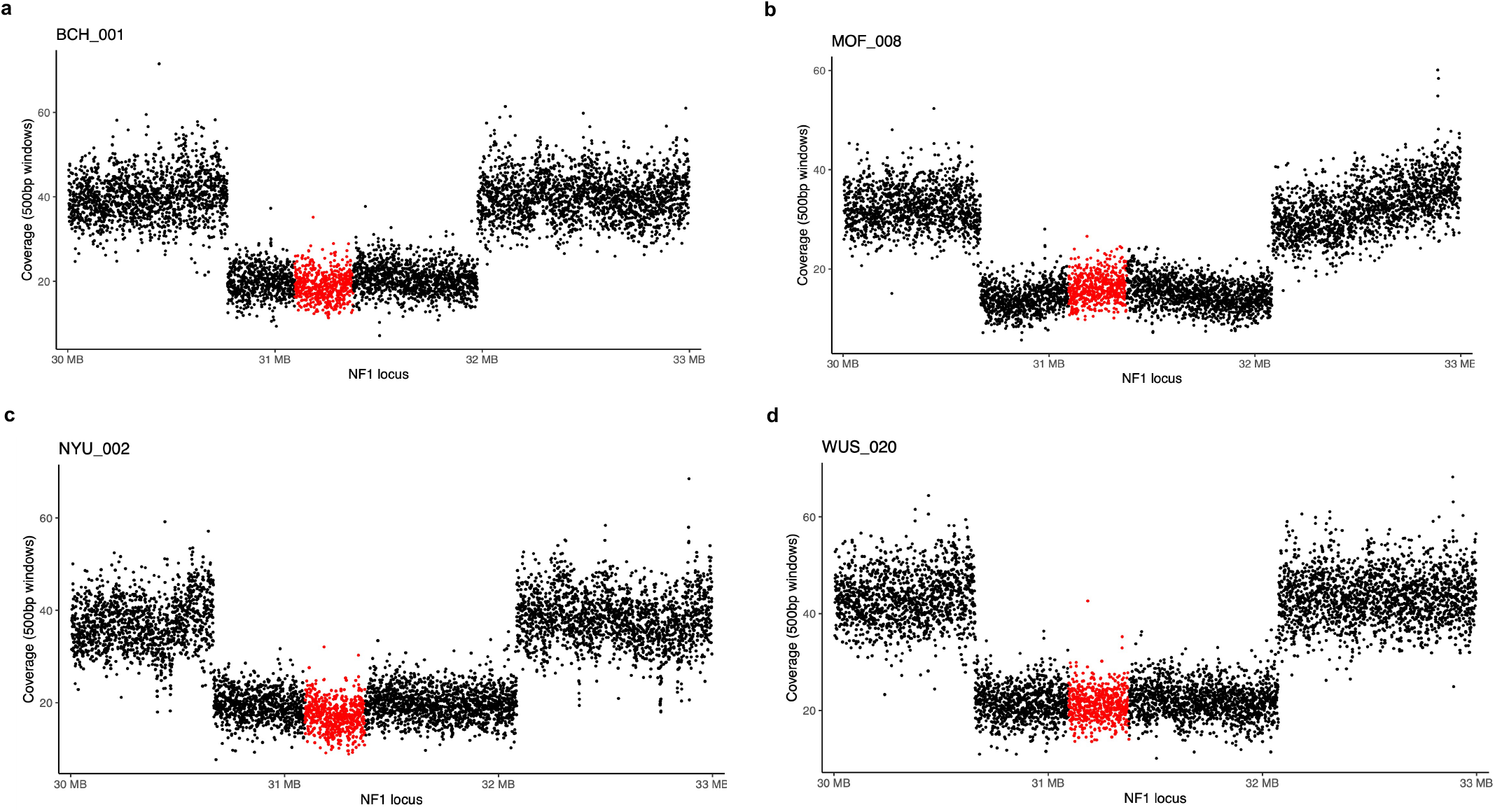
Analysis of germline *NF1* microdeletions. Sequencing coverage calculated at 500 bp windows at the *NF1* locus for individuals BCH_001 (a), MOF_008 (b), NYU_002 (c), and WUS_020 (d). Dots corresponding to windows mapping to *NF1* are shown in red. Genomic coordinates are shown in megabases (MB) according to GRCh38.

**Supplementary Figure 2.**
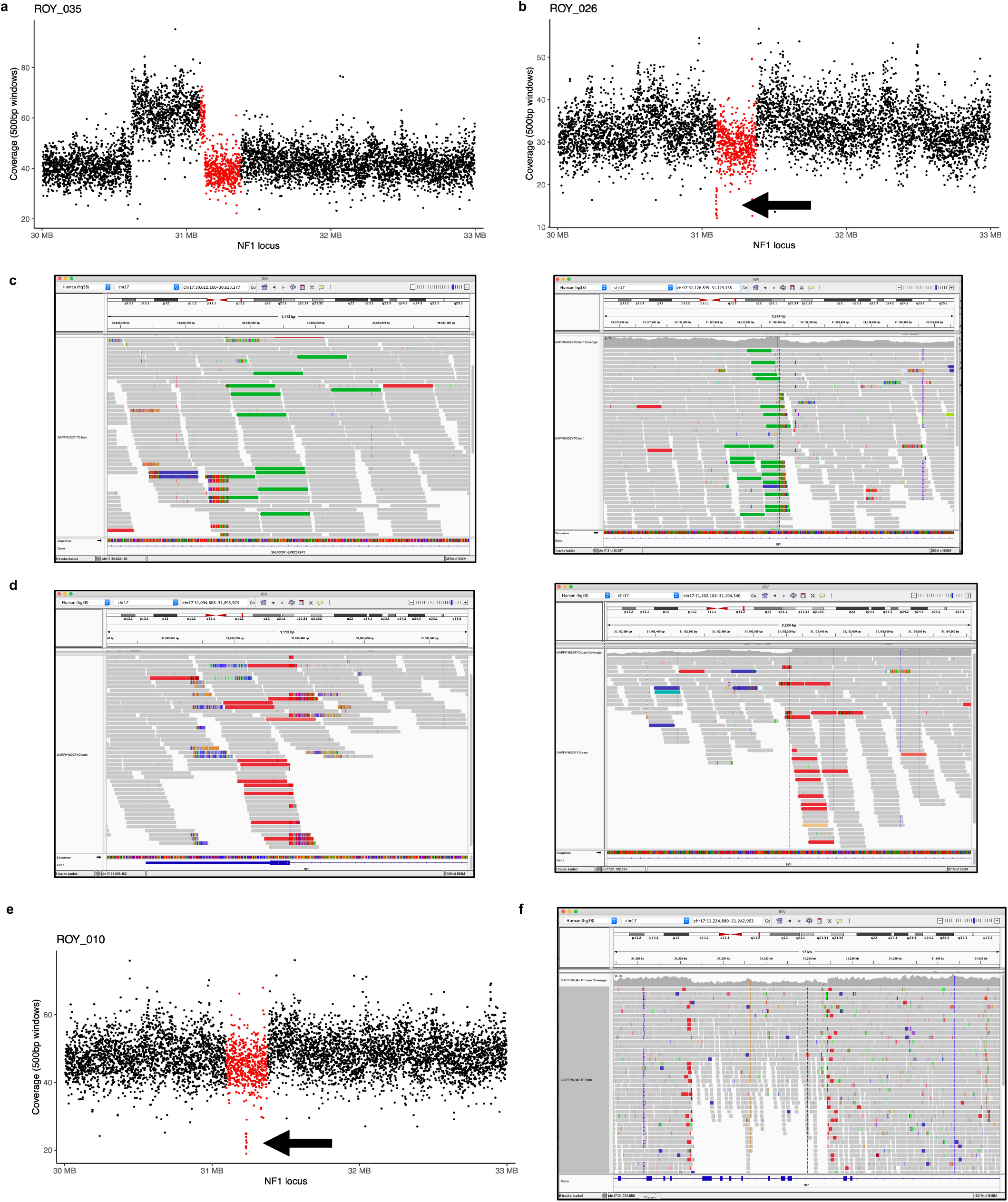
Analysis of germline rearrangements in *NF1*. Sequencing coverage calculated at 500 bp windows at the *NF1* locus for individuals ROY_035 (a) and ROY_026 (b). c, Sequencing reads supporting the tandem duplication detected in ROY_035. d, Sequencing reads supporting the intragenic deletion detected in ROY_026. e, Sequencing coverage calculated at 500 bp windows at the *NF1* locus for individual ROY_010. f, Sequencing reads supporting the intragenic deletion detected in ROY_010. Dots corresponding to windows mapping to *NF1* in a, b, and e, are shown in red, and genomic coordinates are shown in megabases (MB) according to GRCh38. Gray = concordance and colours = discordance with GRCh38 reference in c, d, and f.

**Supplementary Figure 3.**
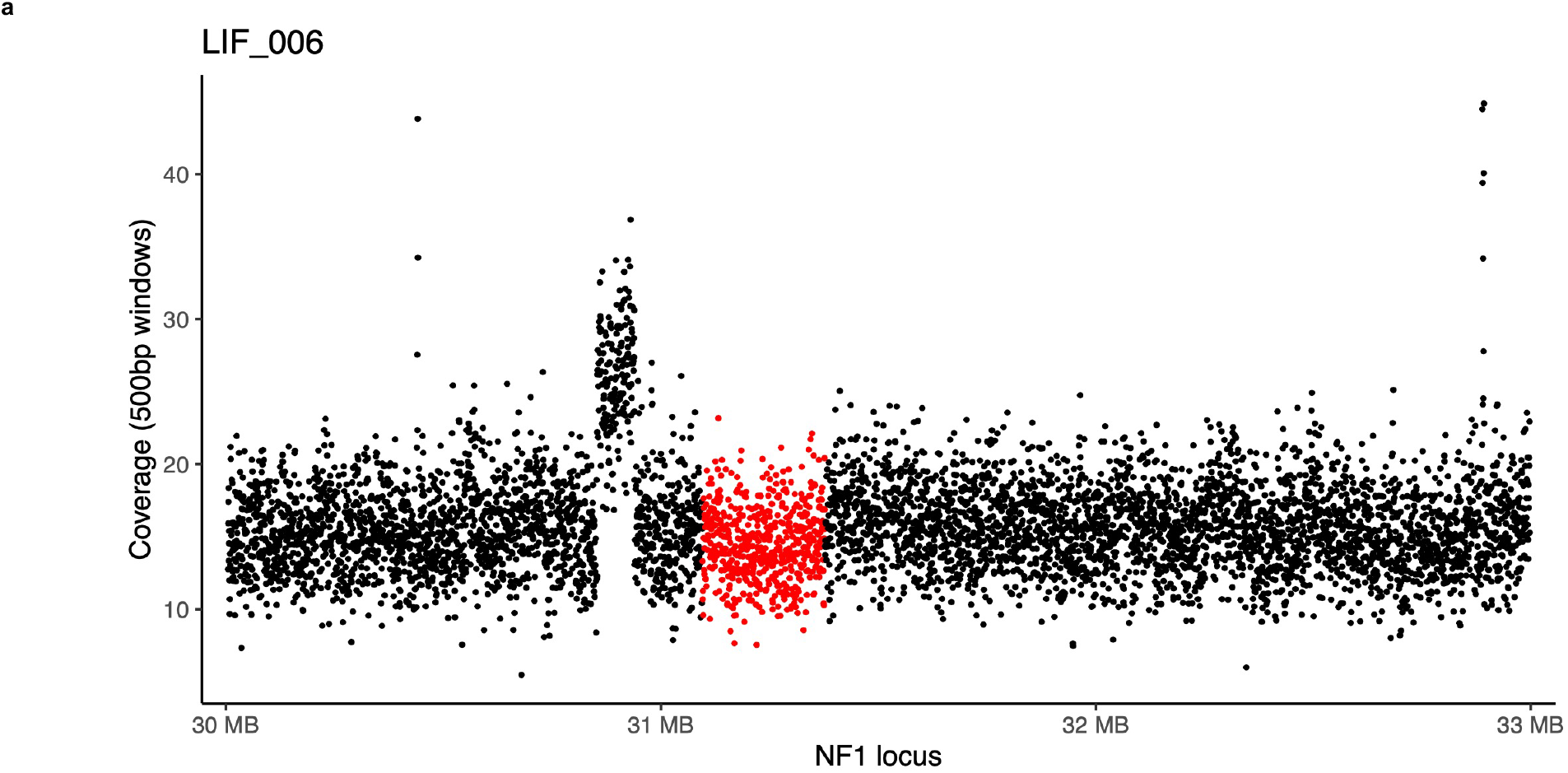
Analysis of germline rearrangements upstream of *NF1.* **a,** Sequencing coverage calculated at 500 bp windows at the *NF1* locus for individual LIF_006. Dots corresponding to windows mapping to *NF1* are shown in red.

**Supplementary Figure 4.**
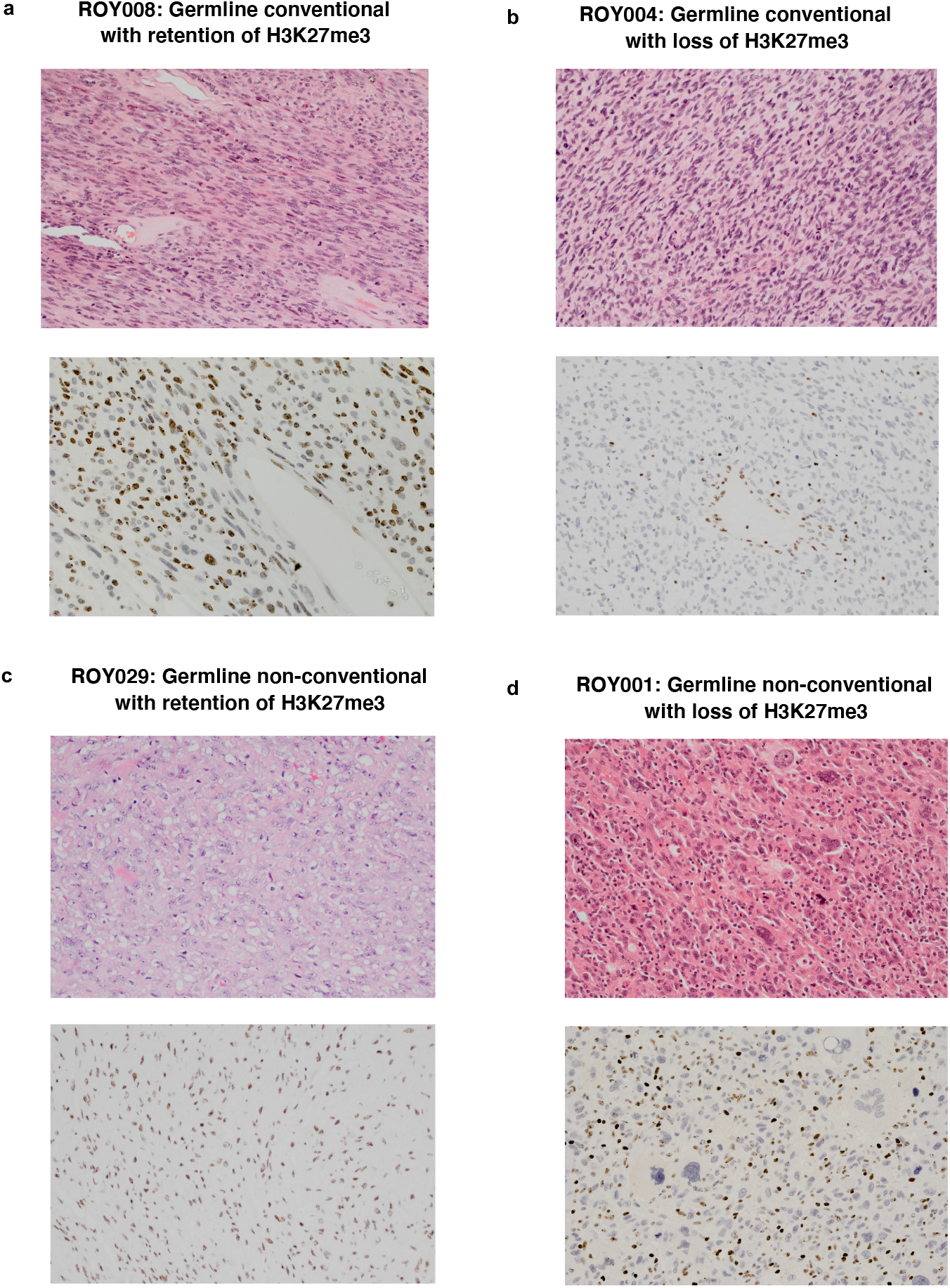
Pathology classification of MPNSTs. H&E and H3K27me3 staining of four representative MPNSTs.

**Supplementary Figure 5.**
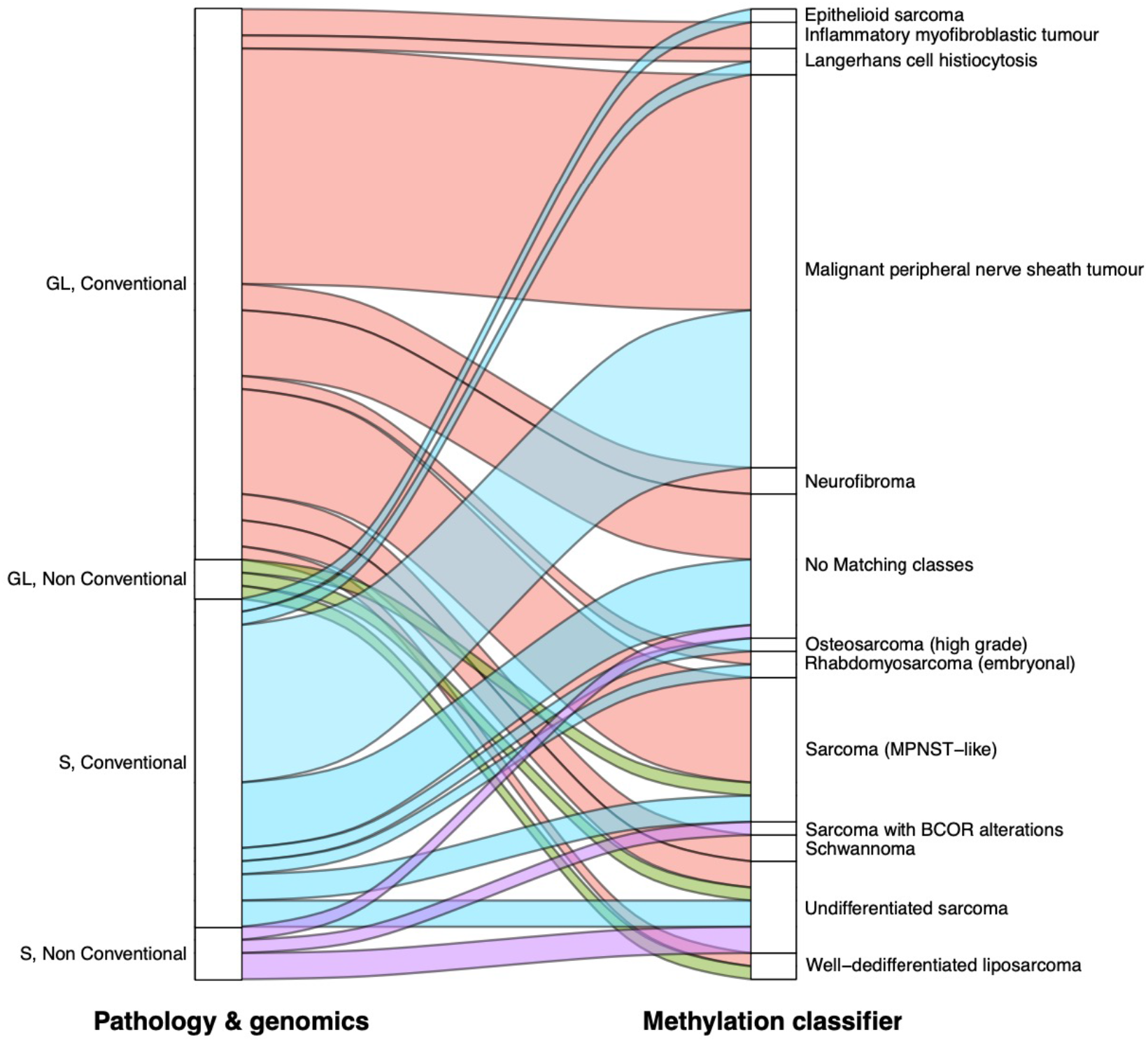
Relationship between the methylation classifier results and the consensus classification established by the GeM consortium based on genomics and pathology information.

**Supplementary Figure 6.**
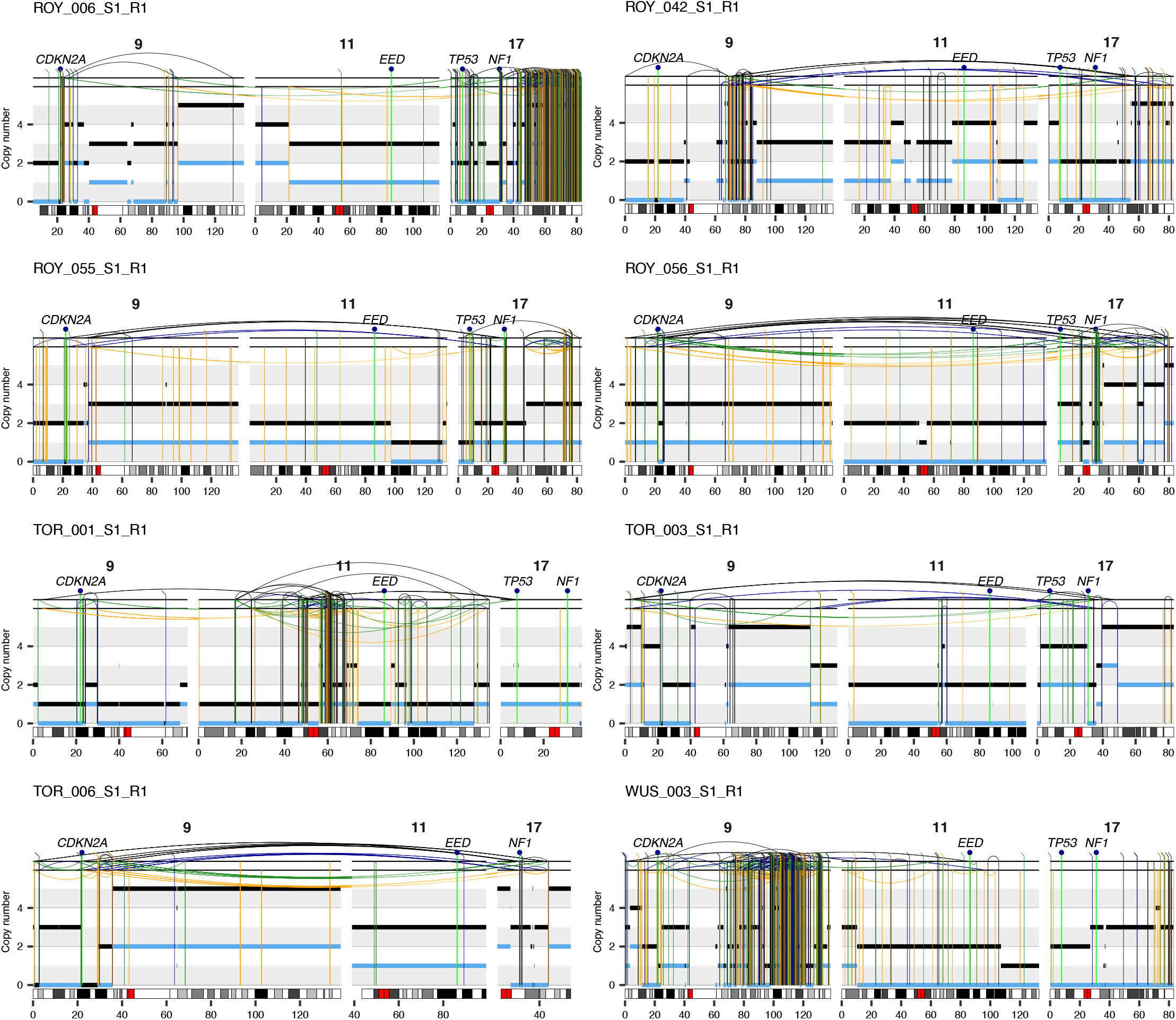
Analysis of complex genome rearrangements involving the *CDKN2A, EED, TP53* and *NF1* loci. Rearrangement profiles of representative MPNSTs showing complex rearrangements of *CDKN2A, EED, TP53* and *NF1.* The position of *CDKN2A, EED, TP53* and *NF1* is shown at the top of each plot. Total and minor copy number values are shown in black and blue, respectively. The chromosome number is indicated on top of each rearrangement profile. Genomic coordinates are shown in Mbp according to GRCh38.

**Supplementary Figure 7.**
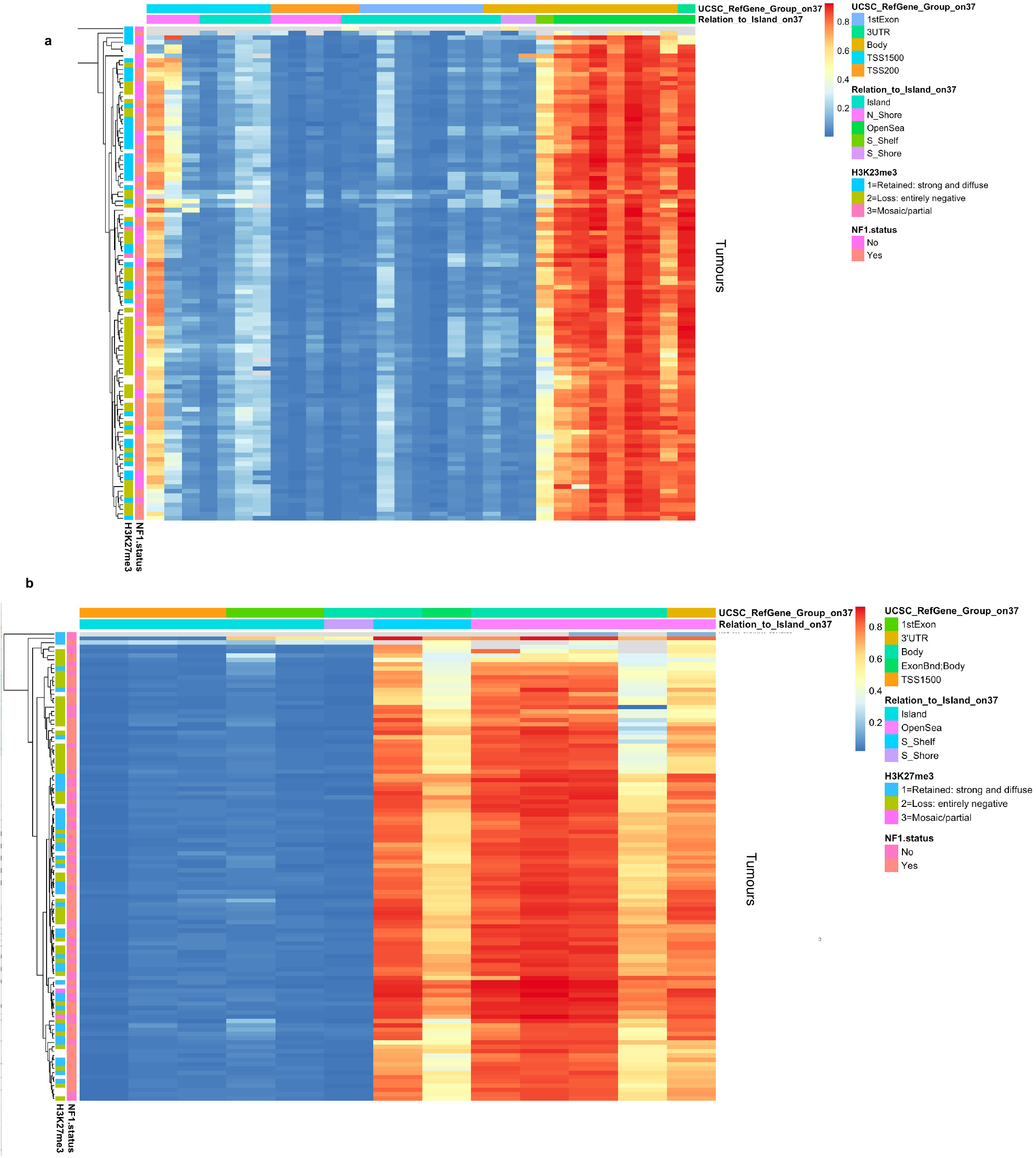
Methylation profiling results for *EED* and *SUZ12*. Beta values for the probes mapping to EED (a) and SUZ12 (b) are shown.

**Supplementary Figure 8.**
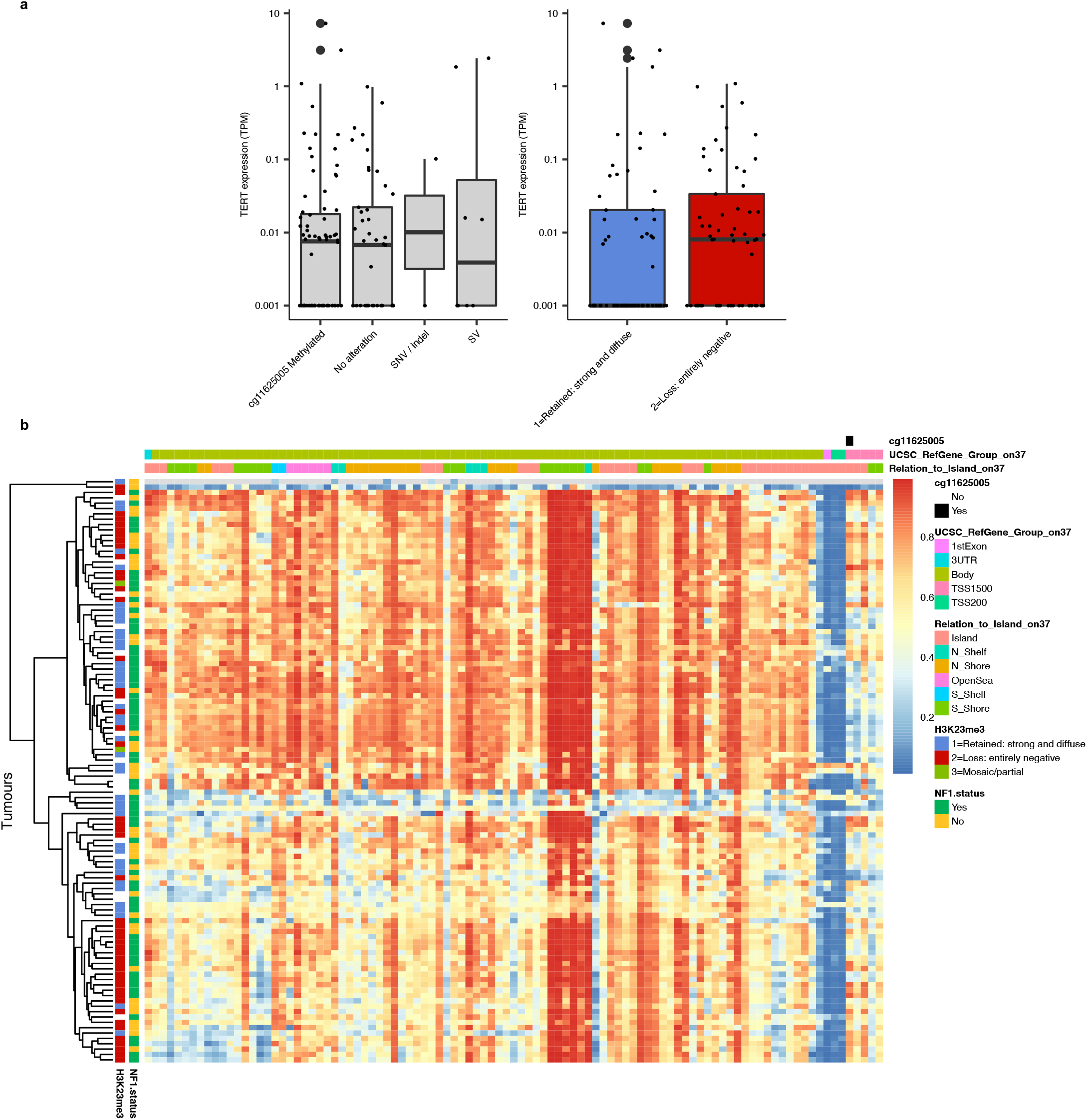
Analysis of replicative immortality in MPNSTs. **a,** *TERT* is overexpressed in samples with *TERT* promoter mutations, SVs mapping to the *TERT* locus (+/- 100Kb), and methylation of the probe cg11625005. b, Beta values for the probes mapping to *TERT*. Each row corresponds to a sample and each column to a probe. The probe cg11625005 is highlighted in black.

**Supplementary Figure 9.**
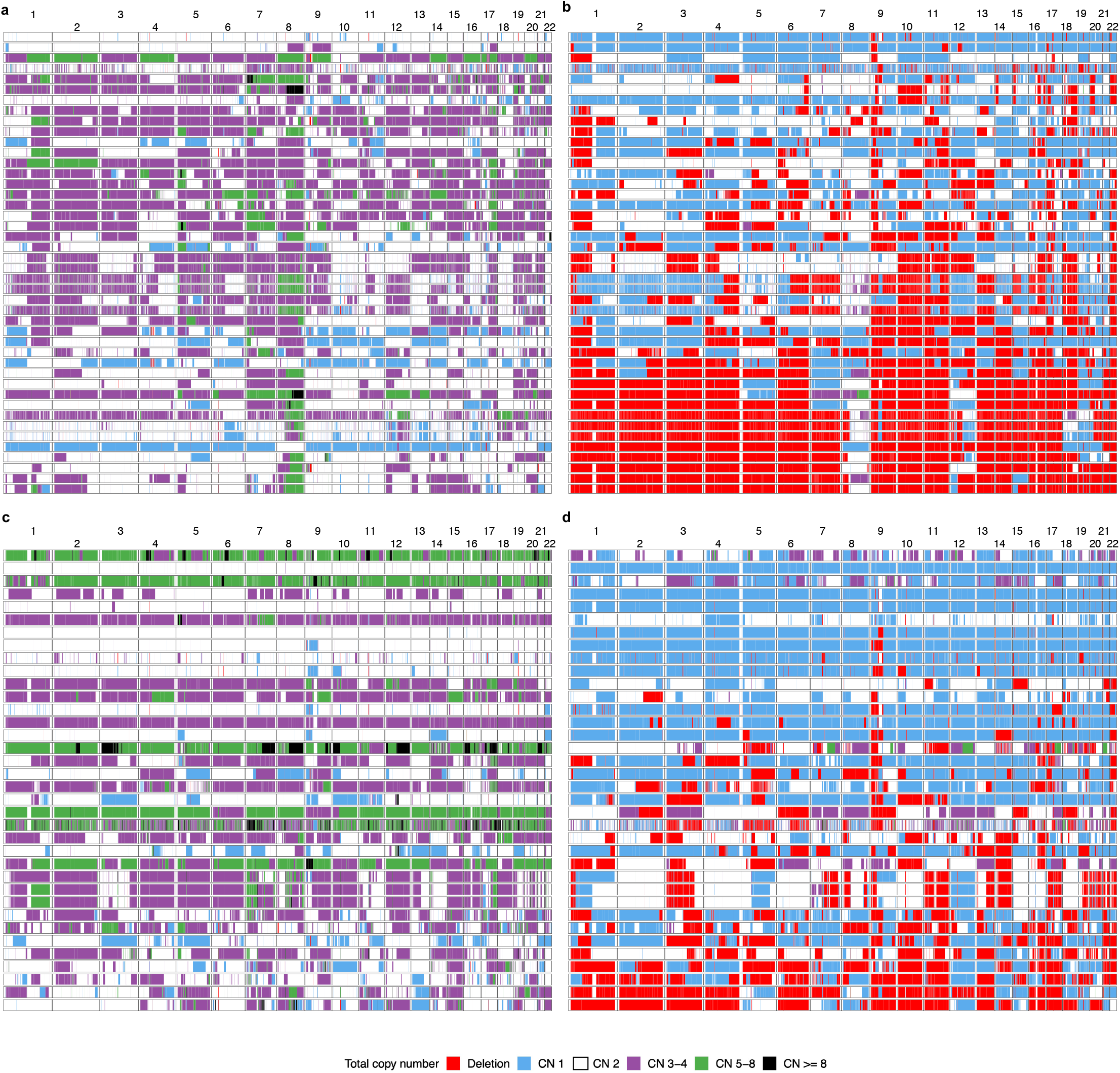
Landscape of somatic copy number aberrations in MPNSTs. Total (a) and minor (b) copy number of the autosomal genome in MPNSTs with loss of H3K27me3. Total (c) and minor (d) copy number of the autosomal genome in MPNSTs with retention of H3K27me3.

**Supplementary Figure 10.**
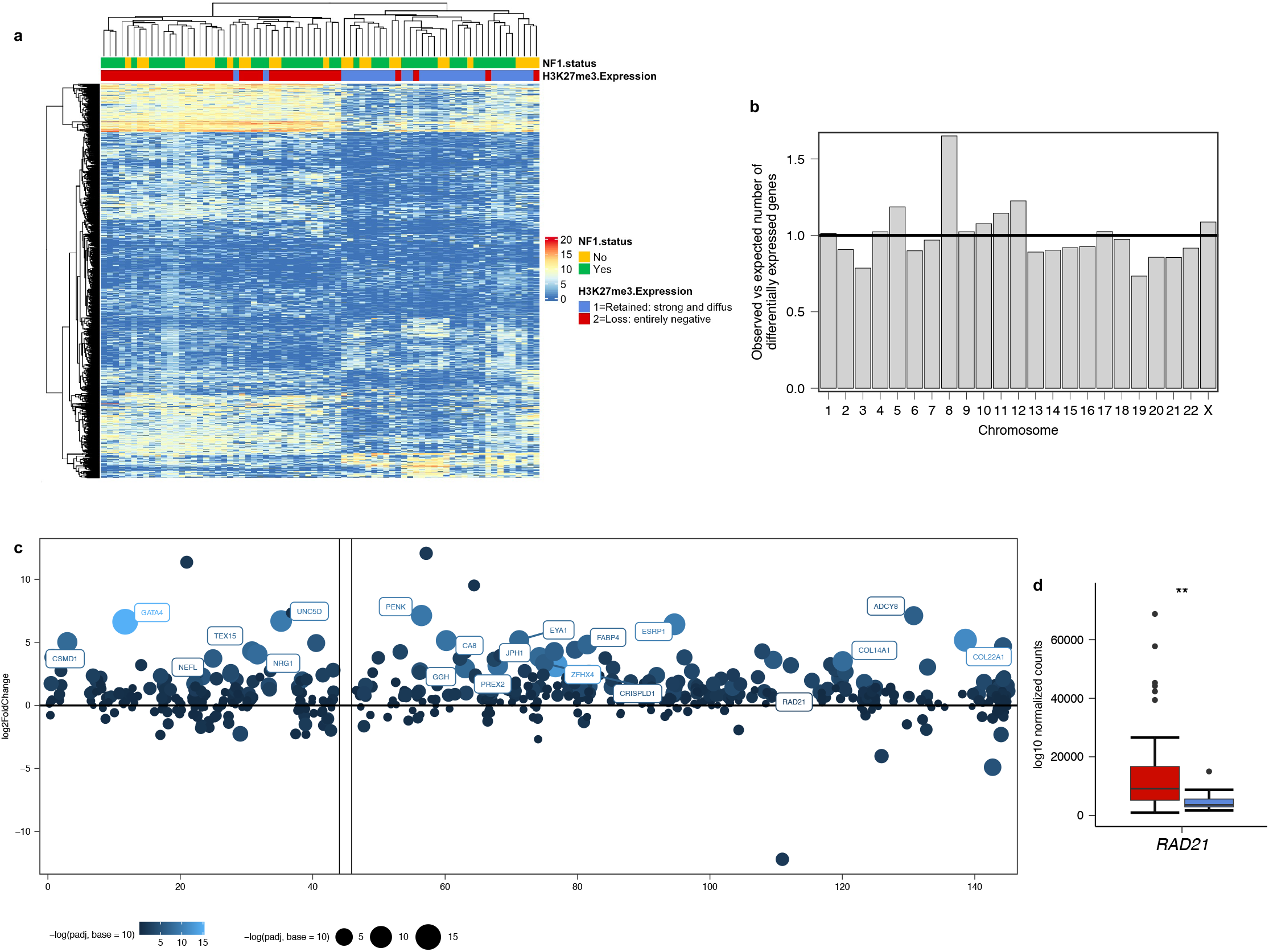
Differential expression analysis. **a,** Heatmap showing the normalised expression counts for genes differentially expressed between MPNSTs with loss or retention of H3K27me3. b, Genes in chromosome 8 are enriched among the set of genes differentially expressed between MPNSTs with loss and retention of H3K27me3. c, Log2 fold change estimate for chromosome 8 genes for tumours with loss of H3K27me3 expression with respect to those with retained H3K27me3. Overall, chromosome 8 genes show higher expression in MPNSTs with loss of H3K27me3. The size and colour of the dots are proportional to the adjusted *P* value calculated in the differential expression analysis. Genomic coordinates are shown in megabases according to GRCh38. d, Differential *RAD21* expression between MPNSTs with loss or retention of H3K27me3. \*\**P* < 0.01.

**Supplementary Figure 11.**
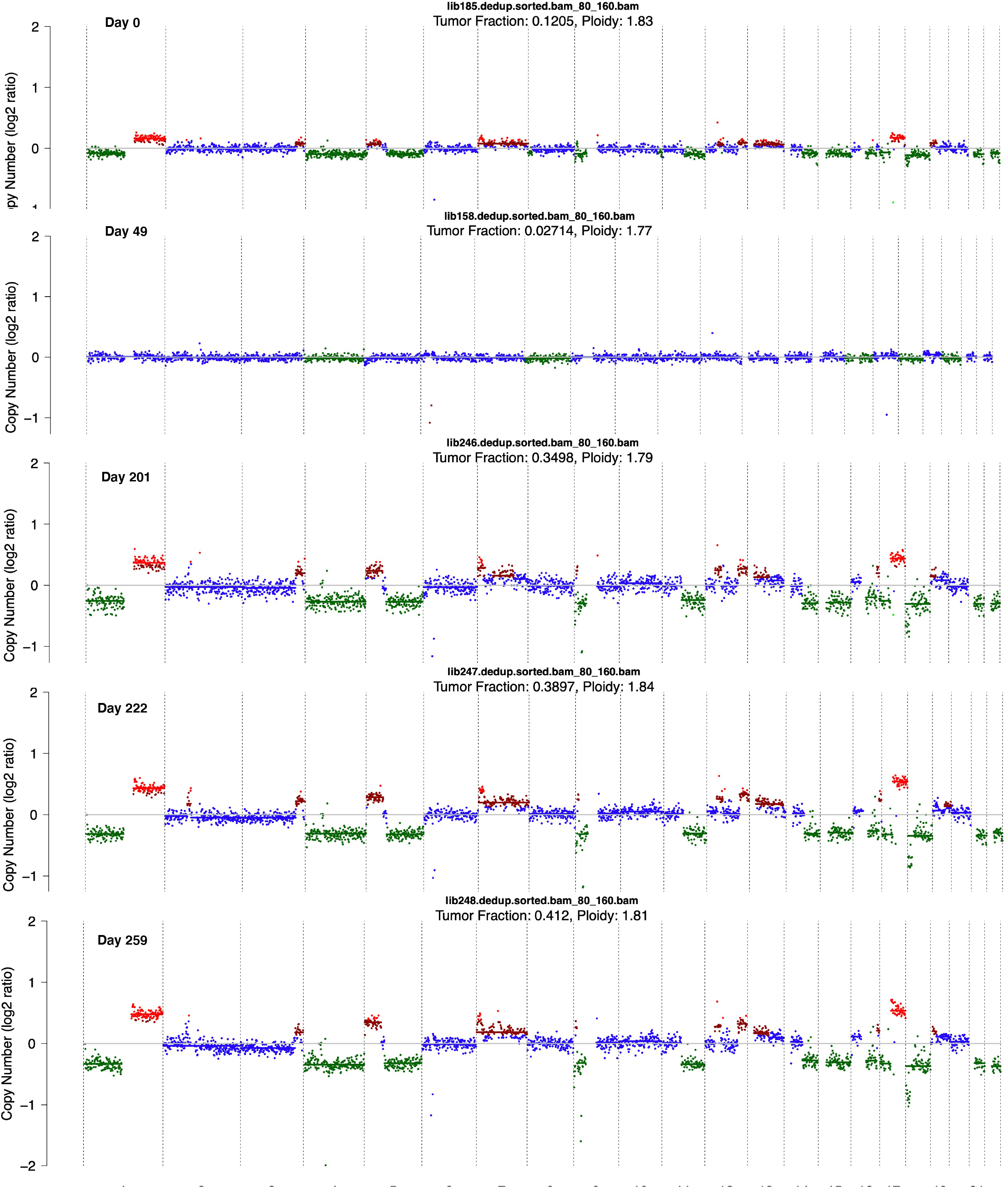
Copy number profiles detected using ultra-low-pass whole-genome sequencing of cell-free DNA. Genome-wide copy number profiles computed using ichorCNA for longitudinal cell-free DNA samples from a patient with MPNST (sar102). The day of collection after clinical diagnosis is indicated in each case. The colour of each dot represents the estimated copy number for the corresponding 1-megabase region. Amplifications are shown in red and deletions in green.

**Supplementary Figure 12.**
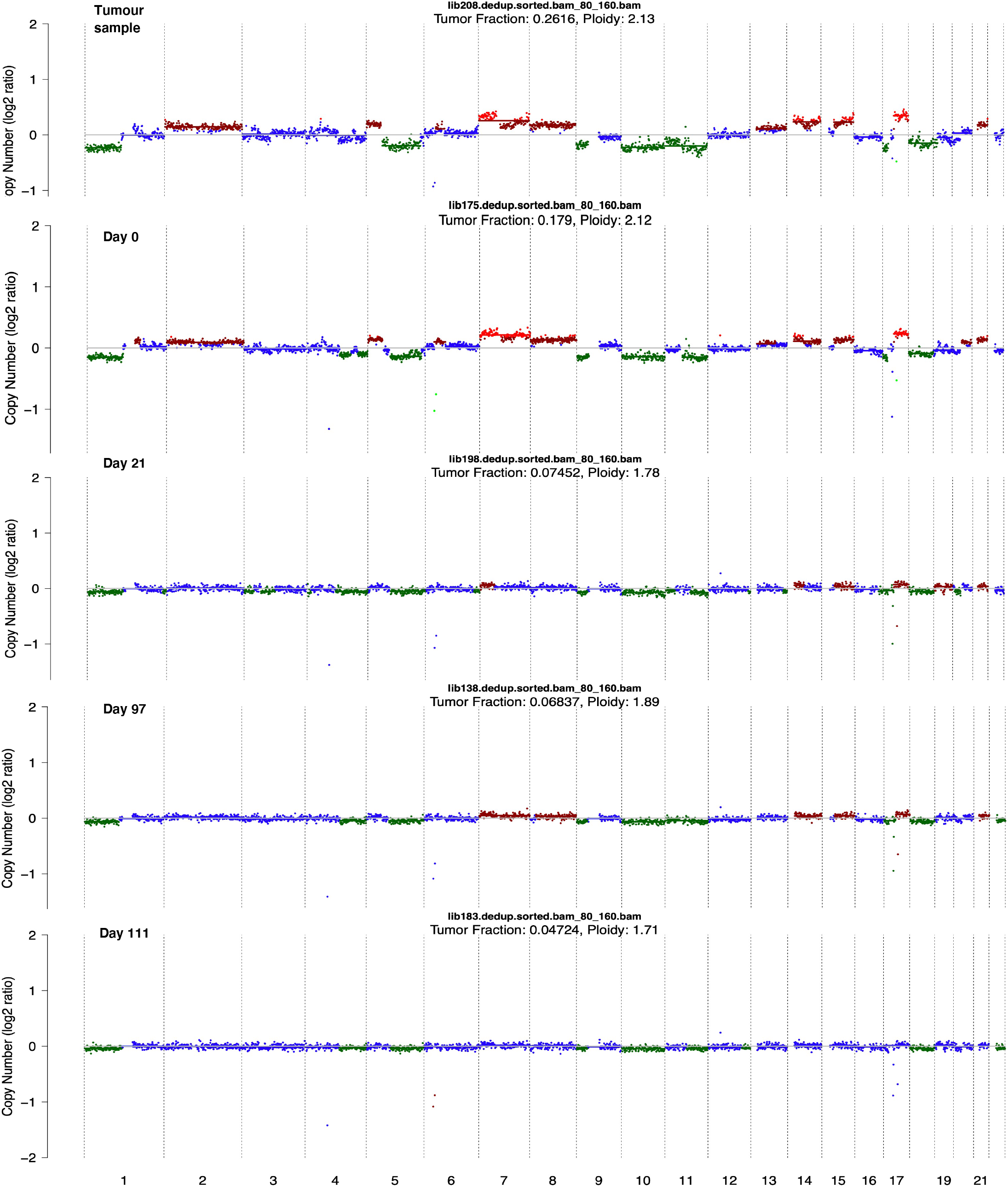
Copy number profiles detected using ultra-low-pass whole-genome sequencing of cell-free DNA. Genome-wide copy number profiles computed using ichorCNA for longitudinal cell-free DNA samples from a patient with MPNST (sar081). The day of collection after clinical diagnosis is indicated in each case. The copy number profile for the tumour sample computed using the same computational and experimental methodology used for cell-free DNA analysis is shown at the top. The colour of each dot represents the estimated copy number for the corresponding 1-megabase region. Amplifications are shown in red and deletions in green.

**Supplementary Figure 13.**
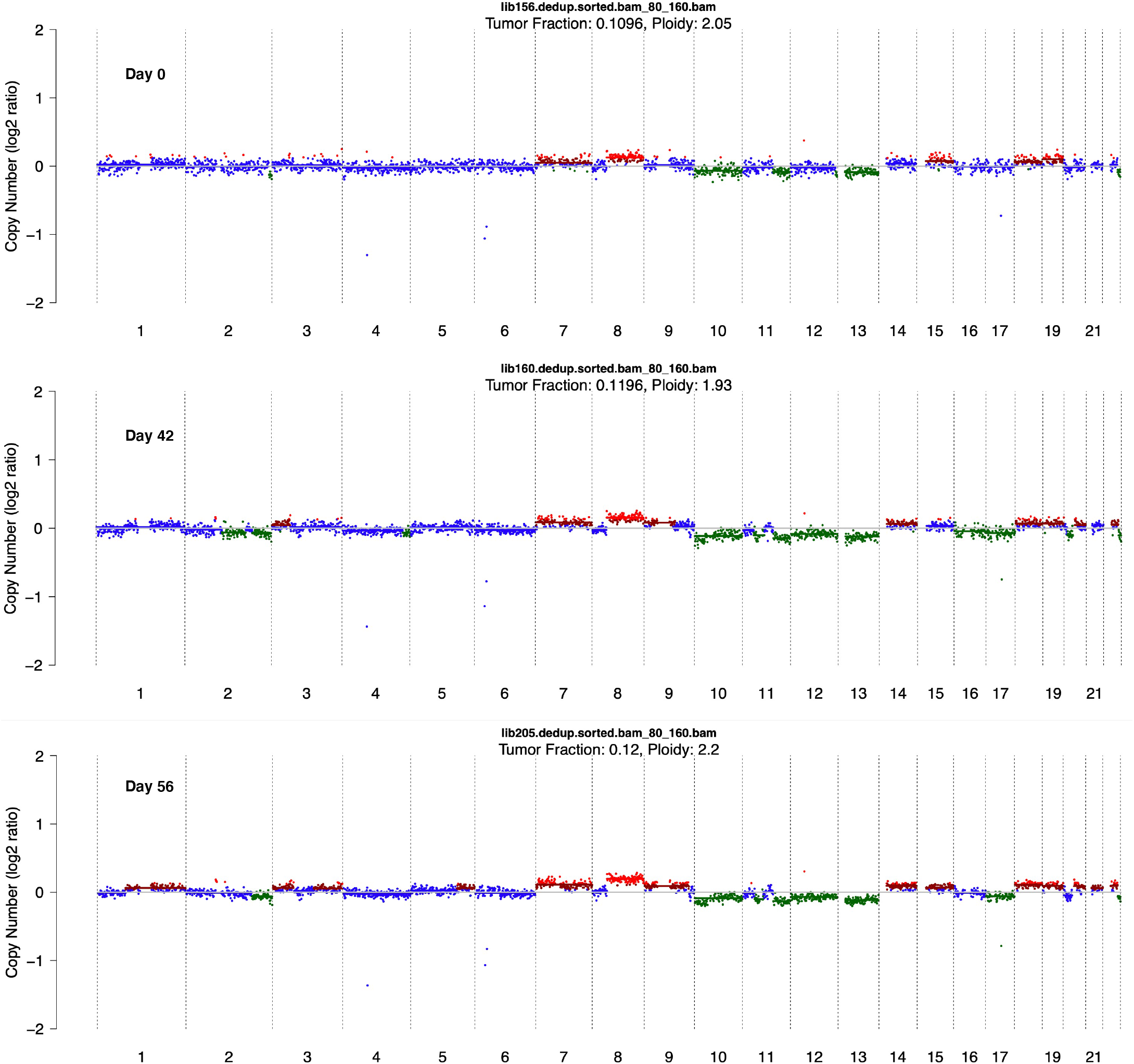
Copy number profiles detected using ultra-low-pass whole-genome sequencing of cell-free DNA. Genome-wide copy number profiles computed using ichorCNA for longitudinal cell-free DNA samples from a patient with MPNST (sar082). The day of collection after clinical diagnosis is indicated in each case. The colour of each dot represents the estimated copy number for the corresponding 1-megabase region. Amplifications are shown in red and deletions in green.

**Supplementary Figure 14.**
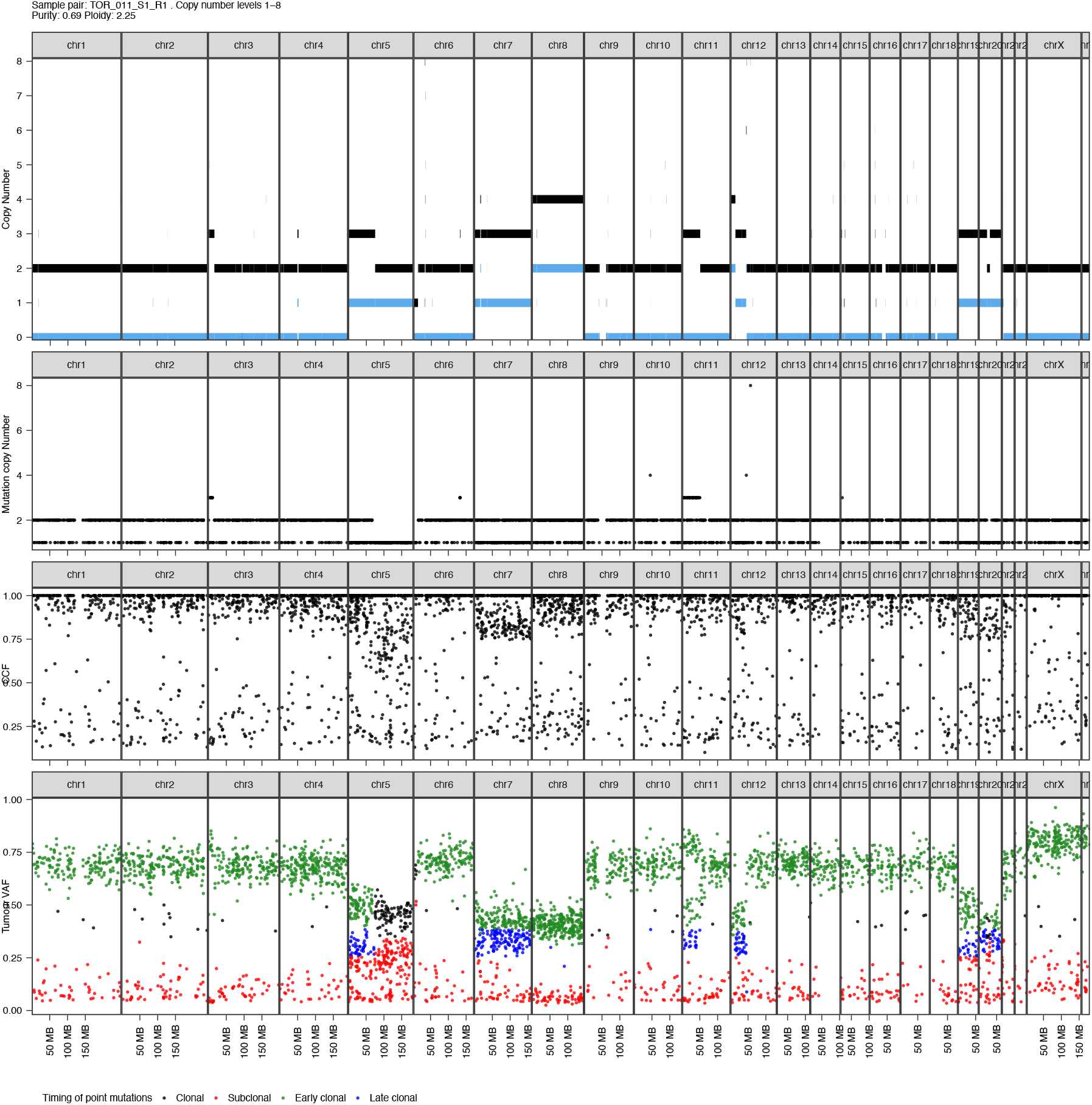
Analysis of the timing of whole genome doubling in MPNSTs with loss of H3K27me3. **a,** Representative example of the somatic copy number profile of an MPNST (TOR_011) with loss of H3K27me3. Total and minor copy number values are shown in black and blue, respectively. **b,** Mutation copy number analysis for SNVs. Each dot represents an SNV and the y axis indicates the number of chromosomal copies in which each mutation is detected in the tumour sample. c, Cancer Cell Fraction (CCF) estimates for each SNV. d, Variant allele fraction (VAF) values in the tumour sample for each SNV coloured according to mutation timing. The large number of clonal SNVs detected in two chromosomal copies in regions with total copy number of 2 and minor copy number of 0 (that is, acquired before whole genome doubling) indicates that the whole genome doubling event occurred late in the clonal evolution of the tumour. Genomic coordinates are shown in megabases according to GRCh38.

**Supplementary Figure 15.**
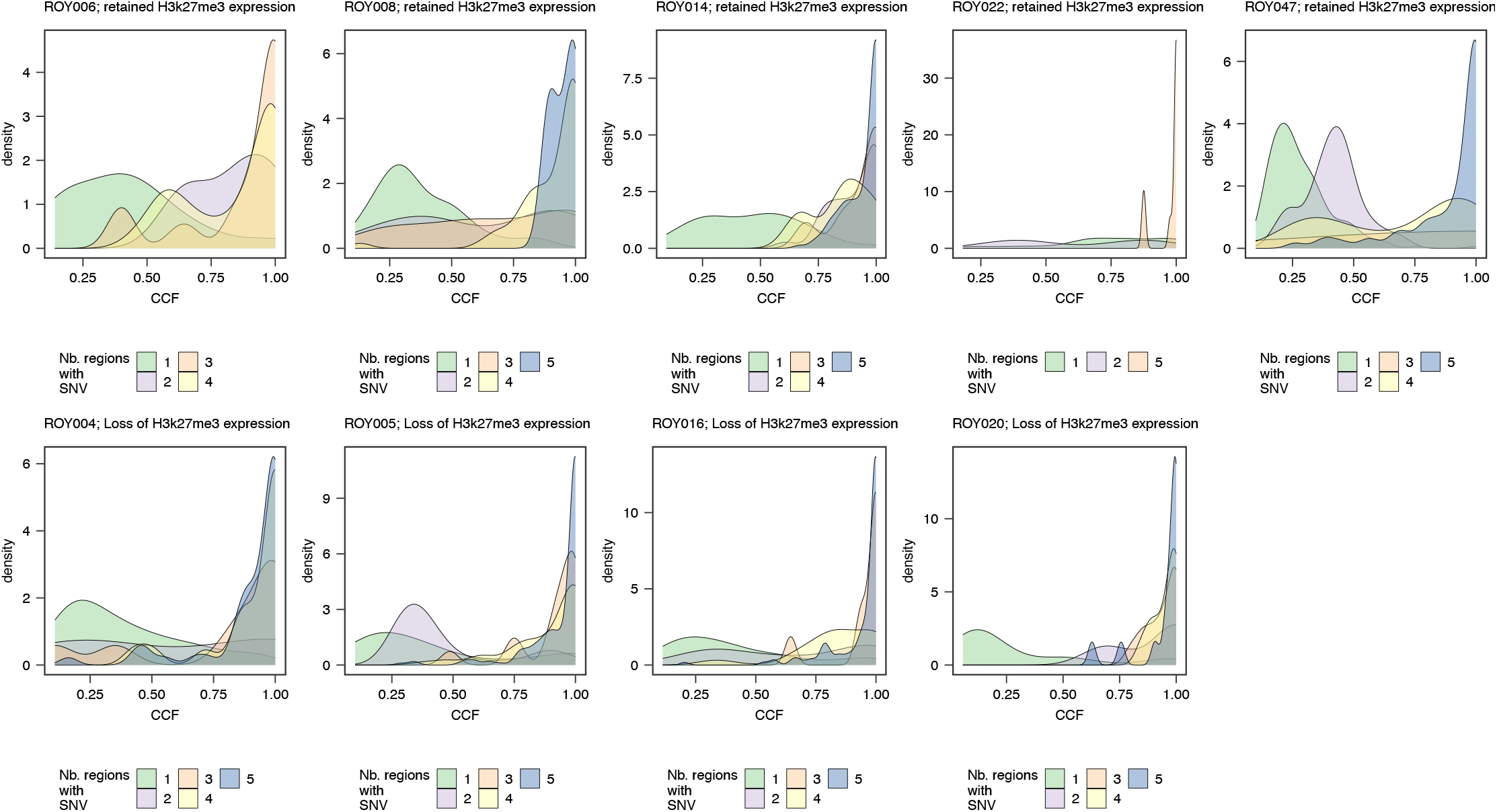
Analysis of the clonality of SNVs detected in the multi-regional exome sequencing data from fresh-frozen samples. The cancer cell fraction (CCF) values for mutations detected in one or multiple regions are shown.

